# Boost or Bust? The impact of supplementation on functional genetic diversity and selective processes in Tasmanian devils

**DOI:** 10.1101/2025.08.03.668351

**Authors:** Andrea L. Schraven, Katherine A. Farquharson, Kimberley C. Batley, Samantha Fox, Andrew V. Lee, Katherine Belov, Luke W. Silver, Carolyn J. Hogg

## Abstract

Translocating individuals into existing populations of conspecifics can support threatened species by increasing population size, maintaining genetic diversity, and reducing the risk of inbreeding. However, for species whose adaptive potential is compromised due to ongoing threats, like disease, the outcome of such management interventions becomes more complex. The Tasmanian devil (*Sarcophilus harrisii*) is a prime example, where the emergence of Devil Facial Tumour Disease (DFTD) has led to significant population declines, raising concerns about their long-term survival. It is therefore critical to understand if the introduction of new functional genetic variants through supplementation actions enhances, or potentially hinders, their long-term persistence. We investigated changes in functional gene diversity at both the population- and individual-levels, pre and post supplementation, across multiple wild devil sites (four supplemented and four not supplemented). We found that functional diversity increased post supplementation. Though the magnitude of change was varied among sites, a similar site-specific pattern was also evident in genome-wide diversity. Importantly, we saw no evidence of swamping of local alleles or those putatively associated with DFTD regressions. This is likely due to the source population representing the broad wild genetic diversity and supplementations facilitating gene flow across the current fragmented landscape. Continued and long-term monitoring at multiple wild sites will be necessary to determine whether future generations retain this introduced genetic variation.

## Introduction

Conservation management interventions could halt the risk of extinction for many threatened species (Gratwicke et al., 2016, Bolam et al., 2021). One management approach is to translocate individuals into a vulnerable population of conspecifics, here referred to as supplementation (also known as reinforcement or augmentation as defined by the IUCN/SSC (2013)). Supplementations enable the introduction of new genetic variation that can diminish the effects of demographic stochasticity and support a population’s adaptive potential (Biebach and Keller, 2009, Johnson et al., 2010, Weeks et al., 2017). Genetic changes in genome-wide diversity following supplementations in threatened populations have been studied (examples include the adder (*Vipera berus*) (Madsen et al., 1999), Florida panther (*Puma pinnatus*) (Johnson et al., 2010), mountain pygmy-possum (*Burramys parvus*) (Weeks et al., 2017), Allegheny woodrat (*Neotoma magister*) (Muller-Girard et al., 2022), and Macquarie perch (*Macquaria australasica*) (Pavlova et al., 2024)), providing evidence that such management interventions can enhance genetic variability in vulnerable populations.

Genome-wide genetic diversity is widely used in conservation management to detect levels of genetic variability at the population- and individual-levels of a species as it is highly influenced by genetic drift, gene flow, and mutation rates (Fernandez-Fournier et al., 2021). However, genome-wide markers have little to no effect on individual fitness (Teixeira and Huber, 2021), and there is growing support for integrating functional genetic data into conservation management investigations (Hoelzel et al., 2019, Theissinger et al., 2023). Variations within the coding (Teixeira and Huber, 2021) regions of the genome that give rise to the breadth of phenotypic differences observed within a population can provide direct insights into the adaptive potential of a species, population, and/or individual (Eizaguirre and Baltazar-Soares, 2014, Hoelzel et al., 2019). Without sufficient genetic variability, populations are highly susceptible to the pressures of inbreeding and reduced fitness (Westemeier et al., 1998, Xue et al., 2015), compromising their long-term viability (Spielman et al., 2004). Where species face strong selective pressures imposed by threatening processes such as climate change, habitat loss, and disease (Palombo, 2021), focussing on functional diversity is useful in determining their genetic resilience and adaptive capacity to changing environments (Lacy, 1997, Kardos et al., 2021). For species suffering from known disease events, common targets for functional investigations include the major histocompatibility complex (MHC) due to its importance in the immune response. For example, alleles of the MHC class II DRB genes have been associated with coronavirus infection susceptibility in *Hipposideros* bat species (Schmid et al., 2023). Many studies of coding regions of the genome typically only investigate a single or a few genes, however, with the increased ease of whole genome sequencing (WGS) or exome sequencing, it is now possible to identify variation across large suites of genes such as those involved in immune function (Farquharson et al., 2022, Silver et al., 2022).

Diseases impose a strong selective pressure on many species and have contributed to both species’ extinctions and significant population declines (De Castro and Bolker, 2005; Smith et al., 2006, Trumbo et al., 2023). Emerging diseases can drive evolutionary responses in affected populations by favouring specific genotypes that confer a resistance and/or tolerance to infection (De Castro and Bolker, 2005, Trumbo et al., 2023). In this context, management strategies that actively increase gene flow must be approached with caution. This is because although these interventions improve genetic diversity and reduce inbreeding they may also inadvertently interfere with natural adaptive processes. A well-documented example is the Tasmanian devil (*Sarcophilus harrisii*), which continues to experience range-wide species’ decline as a result of Devil Facial Tumour Disease (DFTD) (Lazenby et al., 2018). DFTD consists of two independent infectious clonal cancers: DFT1, that was first detected in 1996 (Hawkins et al., 2006) and has spread across Tasmania causing local population declines of more than 80% (Cunningham et al., 2021); and DFT2, that was first detected in 2014 and is currently localised to the south-eastern region of Tasmania (Pye et al., 2016b). Multiple investigations have indicated that devils are showing genetic responses to DFT1. A small number of devils from the northwest have been observed either mounting an immune response (Pye et al., 2016a), or showing signs of tumour regression to DFT1 (Wright et al., 2017). A number of candidate loci have been proposed to be involved in these responses (Pye et al., 2016a, Wright et al., 2017, Margres et al., 2018a. In addition, rapid shifts in allele frequencies at candidate loci linked to possible immune- and cancer-related function have also been observed at three devil sites (Epstein et al., 2016), along with a few loci linked to survival in females following DFT1 infection (Margres et al., 2018a). These findings have prompted concerns that supplementing wild and disease-affected sites with ‘DFTD-naïve’ individuals could potentially swamp out localised adaptations to the disease (Hohenlohe et al., 2019, Hamede et al., 2021). Instead “management of adaptive genetic diversity should be prioritised” to allow natural selection to take place (Hamede et al., 2021). While devils appear to be persevering at low population densities (Lazenby et al., 2018), mostly due to precocial breeding of juvenile females at DFTD-affected sites (Jones et al., 2008, Lazenby et al., 2018), they are also succumbing to small and isolated population pressures that threatens their adaptive potential for future threatening processes (Keller and Waller, 2002, Hoelzel et al., 2019, Cunningham et al., 2021, Farquharson et al., 2022).

In 2003, the Australian and Tasmanian governments responded to the threat of a species-level extinction caused by DFTD, by establishing the Save the Tasmanian Devil Program (STDP) with the aim to ensure a self-sustaining and ecologically functional wild population of Tasmania devils (DPIPWE, 2010). The STDP and its collaborators have produced extensive research output on biological and evolutionary processes between host and disease (Cheng et al., 2012, 2017, Kwon et al., 2020, Pye et al., 2021, Batley et al., 2025); developed and maintained a genetically representative insurance metapopulation (CBSG, 2008, Hogg et al., 2015, 2017, Farquharson et al., 2022), including two disease-free and free ranging populations (Thalmann et al., 2016, Huxtable et al., 2019, Wise et al., 2019); and provided long-term monitoring of multiple wild sites across their range (Lazenby et al., 2018). While effective treatments, or vaccine developments, are still progressing (Pye et al., 2018, 2021), the STDP are managing wild sites in the presence of DFTD. In 2015, the STDP commenced the Wild Devil Recovery (WDR) project to trial supplementations as a potential strategy to support wild populations (Fox and Seddon, 2019). Seven releases to four sites have now occurred, answering several demographic and genetic questions (Fox and Seddon, 2019). Increasing the population size through supplementation at one site had no impact on DFTD prevalence and improved both functional and genome-wide diversity (McLennan et al., 2024), whilst at multiple sites genome-wide diversity was maintained over time following releases (Schraven et al., 2025). Given the selective pressure of DFTD, a similar multi-site investigation is needed to determine whether supplementations using ‘DFTD-naïve’ individuals disrupts potential local adaptations (Hohenlohe et al., 2019, Hamede et al., 2021).

Here, we aimed to determine 1) if supplementations to wild sites increase genetic diversity at functional genes, and 2) whether supplementations impacted existing selective processes (i.e. genetic swamping). To achieve our aims, we compared functional diversity, represented by reproductive, immune (including MHC class I genes) and putatively DFTD-associated loci, at the population-level by comparing “not supplemented” and “supplemented” wild devil sites pre and post supplementation. We also compared this functional diversity at the individual-level pre and post supplementation by comparing “incumbents” (i.e. individuals with two wild parents from that site) and “hybrids” (i.e. individuals with one wild site parent and one release cohort parent, noting this also includes any descendants of these F1 hybrids).

## Methods

### Study Sites

Since 2014, the STDP has conducted annual monitoring of eight wild devil sites: four supplemented (either as one-off releases or multiple releases) and four not supplemented (controls; monitored only) (Figure 1A). *Supplemented sites:* Narawntapu was the first wild site supplemented in 2015 with devils (N = 20) sourced from the captive insurance population (IP) (Hogg et al., 2017). However, due to the high mortality rate of released individuals from vehicle strikes shortly after their release (Grueber et al., 2017), all releases from 2017 onwards sourced devils exclusively from the disease-free population located on Maria Island (MI) (Hogg et al., 2020). One-off releases occurred at Stony Head (in 2016, N = 33) and wukalina (in 2017, N = 33). Multiple releases across several years occurred at Buckland (in 2018, N = 24; in 2020, N = 9) and Narawntapu (in 2019, N = 20; in 2021, N = 10). *Not supplemented sites* were Bronte, Fentonbury, Granville Harbour, and Kempton. Sites selected for supplementation were prioritised based on their low genetic diversity and the projected rate of further diversity loss due to poor connectivity relative to other STDP annual monitoring sites (Grueber et al., 2019). For full details of each devil site included in this study, see Schraven et al. (2025).

**Figure 1.**
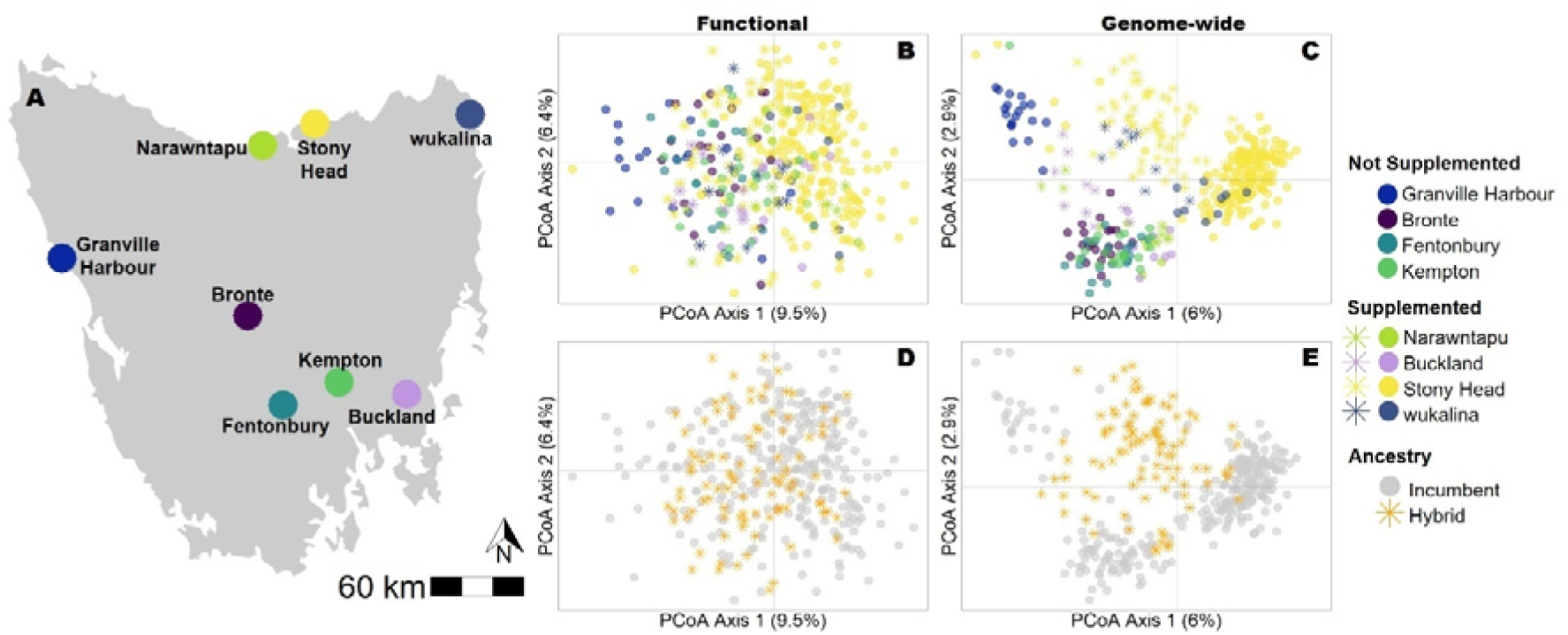
(A) Locations of wild devil sites included in this study across Tasmania, Australia. (B) PCoA of functional diversity at reproductive and immune-related SNPs across eight wild devil sites (N = 325 loci, 379 devils). (C) PCoA of genome-wide diversity across eight wild devil sites (N = 1778 loci, 372 devils). (D) as per (B) but with all incumbent devils shown in grey and all hybrid devils shown orange. (E) as per (C) but with all incumbent devils shown in grey and all hybrid devils shown orange. Sample sizes for each dataset can be found in Table S1.

### Sample Collection and Preparation

Devils were trapped (from 2014 to 2022) during annual monitoring trips led by STDP staff to monitor the health and welfare of individuals and track the prevalence of DFTD. Multi-purpose PVC pipe traps were used (Hawkins et al., 2006), and devils were handled by trained STDP staff under the STDP’s standard operating procedures. DFTD presence and severity was recorded for all trapped individuals (detailed below in *‘DFTD prevalence’*) and an ear biopsy collected in 70% ethanol and stored at −20°C. DNA was extracted from ear biopsies using the MagAttract HMW DNA kit (Qiagen, Germany). The concentration and quality of DNA was assessed using a Nanodrop 2000 Spectrometer (ThermoFisher Scientific) and 0.8% agarose gel electrophoresis at 90 V for 30 min. Full details of sample preparation and assessment are described in McLennan et al. (2019).

### Functional SNPs

A targeted gene sequencing approach was used to assess the functional diversity among and within wild devil sites. We opted for a custom probe design as while whole genome resequencing costs decreased during our study period, this type of short-read sequencing is problematic for devils in that they have a high sequence similarity at their MHC-I genes (>97.7%), a key gene family involved with their ability to respond to DFTD (Cheng et al., 2021). Unless long-read sequencing is used, individual alleles and haplotypes cannot be called with confidence with short-read data (Cheng et al., 2021). Costs associated with long-read whole genome resequencing were prohibitive for this study. As a result, we undertook two different sequencing methods, the first a custom bait for 556 functional genes, including immune (N = 319) and reproductive (N = 237) genes, as well as SNP loci (N = 382) previously associated with selection imposed by DFTD see Farquharson et al. (2022) for details on probe design). A second MHC-I specific assay was also used (Cheng et al., 2021).

The **custom bait** libraries were prepared from extracted DNA using a KAPA HyperPrep Kit (Roche) and following standard procedures. In summary, the DNA mass was standardized to approximately 400 ng and cleaned using a 3x volume of KAPA Hyper-Pure magnetic beads with the concentration of the resulting supernatant measured by Nanodrop and standardised to 120 ng of DNA. The DNA was fragmented at 37°c for 35 min, before A-tailing, end repair, and adapter ligation were performed for each sample. Libraries were cleaned using 0.8x magnetic beads, before amplification. Amplified libraries immediately underwent a 1x bead clean-up before the concentration of each library was determined via Nanodrop and the average library size was estimated on a BioAnalyzer 2100 (Agilent). From this, samples were pooled into groups of 8 with equimolar concentrations for a total of 500 ng of DNA.

Target capture with custom baits (described above) were performed using the myBaits Hybridisation capture for Targeted NGS kit (Daicel Arbor Biosciences) and following standard protocols. Baits were first hybridised for 60°c for 10 min, before blockers were added to the pooled sample libraries. The hybridisation mix was added to the blocker mix containing the pooled libraries and incubated overnight at 65°c then, a bind and wash bead clean-up and library resuspension was performed, libraries were amplified and then cleaned using 1x beads.

To test the efficacy of the target capture protocol, a qPCR was performed on two control regions (GADPH and GUSB) and two enriched regions (DAB and LTA) (see Farquharson et al. (2022) for details on primer and PCR reactions). In summary, regions were amplified using the Quantifast Sybr Green PCR MasterMix (Qiagen), and the copy number threshold was analysed using RotorGene Q Series Software (Qiagen). The enriched libraries were further pooled equimolarly into one final pool, containing 96 samples, and were cleaned using 13x beads. The final library was paired end sequenced on a NovaSeq 6000 SP (Illumina) at Ramaciotti Center for Genomics (UNSW, Sydney, Australia). As 332 samples included in the current study were sequenced as part of prior work (Farquharson et al., 2022, McLennan et al., 2024), two samples were repeated to test the amplification and sequencing accuracy. Additionally, four were removed due to low sequencing quality (N = 2) or a sample mix-up identified by incorrect labelling (N = 2), resulting in 379 devils from eight wild sites (n = 299 at supplemented sites; n = 80 at not supplemented sites).

The sequencing quality of the raw demultiplexed files were assessed using FastQC (Andrews, 2010) and MultiQC (Ewels et al., 2016). Once adapters were trimmed using ‘trimmomatic’ v 0.38 (Bolger et al., 2014), the sample reads from multiple lanes were merged. Reads were then aligned to the Tasmanian devil reference genome (RefSeq mSarHar1.11, GCF_902635505.1) using BWA v 0.7.17 (Li and Durbin, 2009), with the resulting reads sorted, indexed, and duplicates marked using Picard v 2.18.4 (Broad Institute Github Repository, 2019) and SAMtools v 0.1.19 (Li et al., 2009).

Variants were called and filtered using GATK v 4.1.9.0 (McKenna et al., 2010), following methods described in Farquharson et al. (2022). In summary, only variants within the target regions were retained using HaplotypeCaller, before a catalogue of variants across all samples were built using GenomicsDBImport. A single variant file containing all samples was then generated using GenotypeGVCF, and only SNP variants were retained using SelectVariants. SNPs were then filtered using VariantFiltration and the following filters: DP (coverage) < 7,740, QD (quality score normalised by allele depth) < 2.0, SOR (strand bias estimated be the symmetric odds ratio test) > 3.0, MQ (mapping quality) < 40.0, MQRankSum (mapping quality rank sum test) < −12.5, and ReadPosRankSum (rank sum for relative positioning of REF versus ALT alleles within reads) < −8.0.

Further filtering of bi-allelic SNPs was carried out in R v 4.4.3 (R Core Team, 2023) with a MAF (minor allele frequency) > 0.01 and heterozygosity ≤ 90%. For the population genetic analysis, SNPs were retained with a LD (linkage disequilibrium) < 0.3 using the ‘SNPRelate’ v 1.33.2 R package (Zheng et al., 2012). Putative DFTD-associated loci were excluded from population genetic analysis as any functional role has not yet been established, retaining only immune and reproductive genes. For haplotype phasing, no LD filter was applied. Reproducibility between replicate sample pairs was calculated and the sample with the least missing data retained for further analysis. Haplotype phasing was conducted with PHASE v 2.1.1 (Stephens et al., 2001, Stephens and Scheet, 2005) for the reproductive and immune genes with at least one SNP variant and a minimum haplotype count of 2. After filtering, 325 bi-allelic SNPs were retained in the linkage pruned dataset and 925 SNPs retained in the non-linkage pruned dataset. Mean error rate at the four replicate pairs was 1.05% SD = 0.52). For population genetic analysis of functional SNPs, a subset of 260 SNPs was used, which excluded the putative DFTD-associated loci. Haplotype phasing resulted in 286 haplotypes across 105 genes (mean = 2.724 haplotypes per gene; range 2-9; SD = 1.383).

The three classical devil **MHC-I** loci were genotyped (N = 346 individuals) following a long-read sequencing protocol described in Cheng et al. (2021). In summary, amplification of the MHC-I loci involved a 3-step PCR method. The three loci were amplified using previously designed forward (/5AmMC6/gcagtcgaacatgtagctgactcaggtcacGTGTCCCCCCCTCCGTCTCAG) and reverse (/5AmMC6/tggatcacttgtgcaagcatcacatcgtagCCTAACTCCCCCTGCTCCTTCTG) primers and the Platinum SuperFi II PCR Master Mix (Invitrogen). Barcoded Universal F/R Primers Plate-96v2 (Pacific Biosciences) were attached to each sample using the Phusion Hot Start II High Fidelity PCR Master Mix (Thermo Scientific). Finally, samples were pooled equimolarly and cleaned using 0.6 AMPure PB magnetic beads (Pacific Biosciences). Libraries were prepared using the SMRTbell Express Template Prep Kit 2.0 (Pacific Biosciences) and sequenced on a PacBio Sequel II platform at the Australian Genome Research Facility. For details on PCR reactions and conditions, see Cheng et al. (2021).

All raw data processing was completed using the PacBio Secondary Analysis Tools on BioConda Grüning et al., 2018 (available at https://github.com/PacificBiosciences/pbbioconda), and following the methods described by Cheng et al. (2021) and Batley et al. (2025). Briefly, ccs v 6.4.0 was used to generate the ccs reads by-strand, before filtering to remove reads with ≤ 5 read passes and ≥ 0.995 read quality. Samples were demultiplexed using lima v 2.7.1, with concatemer and partial reads removed using isoseq refine v 3.8.2. The resulting reads were aligned to the devil MHC-I UA reference sequence using pbalign v 0.4.1, and samples with less than 250 ccs reads remaining were removed. Alleles were called against a database of known devil MHC-I alleles using Bellerophon (https://github.com/yuanyuan929/bellerophon), and grouped into previously defined supertypes (Cheng et al., 2021).

### Genome-wide SNPs

For genome-wide diversity analysis, we included all 379 individuals from the target capture approach above. Samples had been previously sequenced using DArTseq^TM^ (Diversity Arrays Technology PL, Canberra; DArT) (Jaccoud et al., 2001) and SNPs were filtered using the ‘dartR’ v 2.9.7 package in R (Gruber et al., 2018) in R v 4.3.1 (R Core Team, 2023), detailed in Schraven et al. (2025). Two individuals from the functional dataset were excluded from the genome-wide dataset as they did not pass DArTseq quality controls, and an additional three individuals removed during ‘dartR’ filtering steps. A total of 1778 loci were retained across 372 individuals.

Pedigrees were reconstructed for supplemented sites (n = 4) to assign individuals to parental categories (incumbents or hybrids as defined in the introduction). Pedigree reconstruction was performed using ‘sequoia’ v 2.11.2 R package (Huisman, 2017), which was confirmed with the triadic maximum likelihood estimator (TrioML; Wang, 2007) in COANCESTRY v 1.0 (Wang, 2011) and a principal coordinate analyses (PCoA) using the ‘adegenet’ R package v 2.1.10 (Jombart, 2008). TrioML was used to calculate relatedness among individuals within sites with results divided by two to obtain mean kinship (MK) estimates, which is the average kinship of an individual relative to all other individuals in the site (including self) (Hogg et al., 2019). We used PCoAs for individuals that could not be assigned a parental category. Here, we visualised the admixture within each supplemented sites and individual ancestry was inferred by which parental group they clustered with. The sample sizes of individual ancestry at supplemented sites can be found in Table S1.

### Population genetic analysis

For all genetic analysis (except for genetic differentiation), we grouped individuals based on supplementation history due to the variable sample sizes at each site. For not supplemented sites, all individuals were pooled across all trap years. For supplemented sites, individuals were grouped into pre supplementation (trapped in years before their first release and including first release year) and post supplementation (trapped in years after release). For Narawntapu and Buckland, which underwent multiple supplementation events, any individual trapped after the first release was classified as post supplementation. We further categorized post supplementation individuals into incumbents and hybrids based on their reconstructed pedigrees.

PCoAs were used to visualise genetic differentiation for both the functional and genome-wide genetic diversity using the ‘adegenet’ R package v 2.1.10 (Jombart, 2008), and also quantified using pairwise F_ST_ implemented in the ‘StAMPP’ R package v 1.6.3 (Wright, 1949, Weir and Cockerham, 1984, Pembleton et al., 2013), with 95% confidence intervals calculated via 2000 bootstraps across loci. Individual genetic diversity using standardised heterozygosity (H_S_) was calculated with the ‘genhet’ R function (Coulon, 2010), where individual observed heterozygosity was divided by the mean observed heterozygosity across all individuals in the functional and genome-wide datasets. Individual inbreeding coefficients (FH) were calculated using PLINK v 1.9 (Chang et al., 2015), and the average FH and associated SE values were calculated for each devil sites and group for both functional and genome-wide datasets.

### Allele frequencies

Immune genes were categorised into eight groups: antimicrobial peptides (AMP); cluster of differentiation (CD); complement genes; cytokines; immunoglobulins (IG), natural killer cells (NK), and toll-like receptors (TLR). The reproductive genes were grouped all together. MHC-I genes were grouped into UA, UB, and UC, and additionally, the frequency of whole gene deletions was analysed. Additionally for MHC, the average number of alleles present in each group (not supplemented, pre and post supplementation incumbents and post supplementation hybrids) was calculated and visualised. The allele frequencies for each gene for each study site were calculated in R v 4.3.1 (R Core Team, 2023). For supplemented sites, we also calculated the allele frequency pre and post supplementation, and between incumbents and hybrids. The differences in allele frequencies were visualised with heatmaps.

### DFTD-associated loci

We examined all allele frequencies of putative DFTD-associated loci that had been previously identified (Epstein et al., 2016, Wright et al., 2017, Margres et al., 2018a, Margres et al., 2018b). Of the 382 DFTD-associated loci that were included in the probe design (Farquharson et al., 2022), only 65 were retained post R filtering. To address potential alterations to local adaptations of these genes, we decided to examine all allele frequencies of DFTD loci (n = 382) previously found to have associations with some aspect of DFTD tolerance and/or resistance (Epstein et al., 2016, Wright et al., 2017, Margres et al., 2018a). As these loci were aligned to the previous reference genome produced by Murchison et al. (2012), we first located their positions on the newer chromosome-level genome (Stammnitz et al., 2023) to investigate the potential function of all 382 loci. We determined their genomic locations (exonic, intronic, etc.) using annovar v 20180416 (Wang et al., 2010) in the devil reference genome (GCF_902635505.1_mSarHar1.11). Substantial shifts in allele frequencies pre and post supplementation (N = 21 loci) were visualised with heatmaps.

### DFTD prevalence

Every individual devil trapped between 2014 and 2022 was scored from one to five for gross DTFD lesions, with a score of four or five considered active DFTD. Prevalence of DFTD was calculated as the proportion of unique trapped individuals with a score of 4 or 5 in each year for each site. Juvenile devils (classified as devils < two years of age) are less likely to be affected by DFTD than mature devils (classified as devils ≥ two years of age) (Cheng et al., 2017). Therefore, we recalculated DFTD prevalence for each trap year at each site again but excluded one-year-old devils.

## Results

### Changes to Diversity

We found limited genetic differentiation at the functional genes among our eight study sites across Tasmania (Figure 1B). In contrast, genetic structure was more pronounced for genome-wide diversity, where the genetic differentiation appeared to reflect the geographic distribution of wild devil sites (Figure 1A and 1C, Supplementary Table S2). Hybrid individuals (one wild parent, one released parent) represented a genetic mixture of both the current wild functional and genome-wide diversity (Figures 1D and 1E). These hybrid individuals appeared to increase the genome-wide similarity among sites (Figure 1E).

There was little difference in the patterns of standardised heterozygosity (H_S_) between functional and genome-wide diversity at both supplemented and not supplemented sites (Figure 2). For not supplemented sites, Granville Harbour had the lowest H_S_ and highest FH values while the other three sites (Bronte, Fentonbury, and Kempton) were relatively similar to each other (all had above the average H_S_ and below the average FH). At the supplemented sites (Buckland, Narawntapu, Stony Head, and wukalina) changes to H_S_ and FH pre and post supplementation were variable, though on average hybrids had greater above average functional and genome-wide diversity than incumbents (Figure 2A & B). Without supplementation to these sites (measured through post supplementation incumbent individuals), we found that H_S_ at functional genes decreased substantially at both Buckland and wukalina while there was a slight increase at Stony Head. Similarly, H_S_ at genome-wide SNPs declined at Buckland in incumbents post supplementation, though increased marginally at Stony Head and wukalina. A comparison of pre and post supplementation could not be made at Narawntapu as no “pre supplementation incumbent individuals” were available in our datasets.

**Figure 2.**
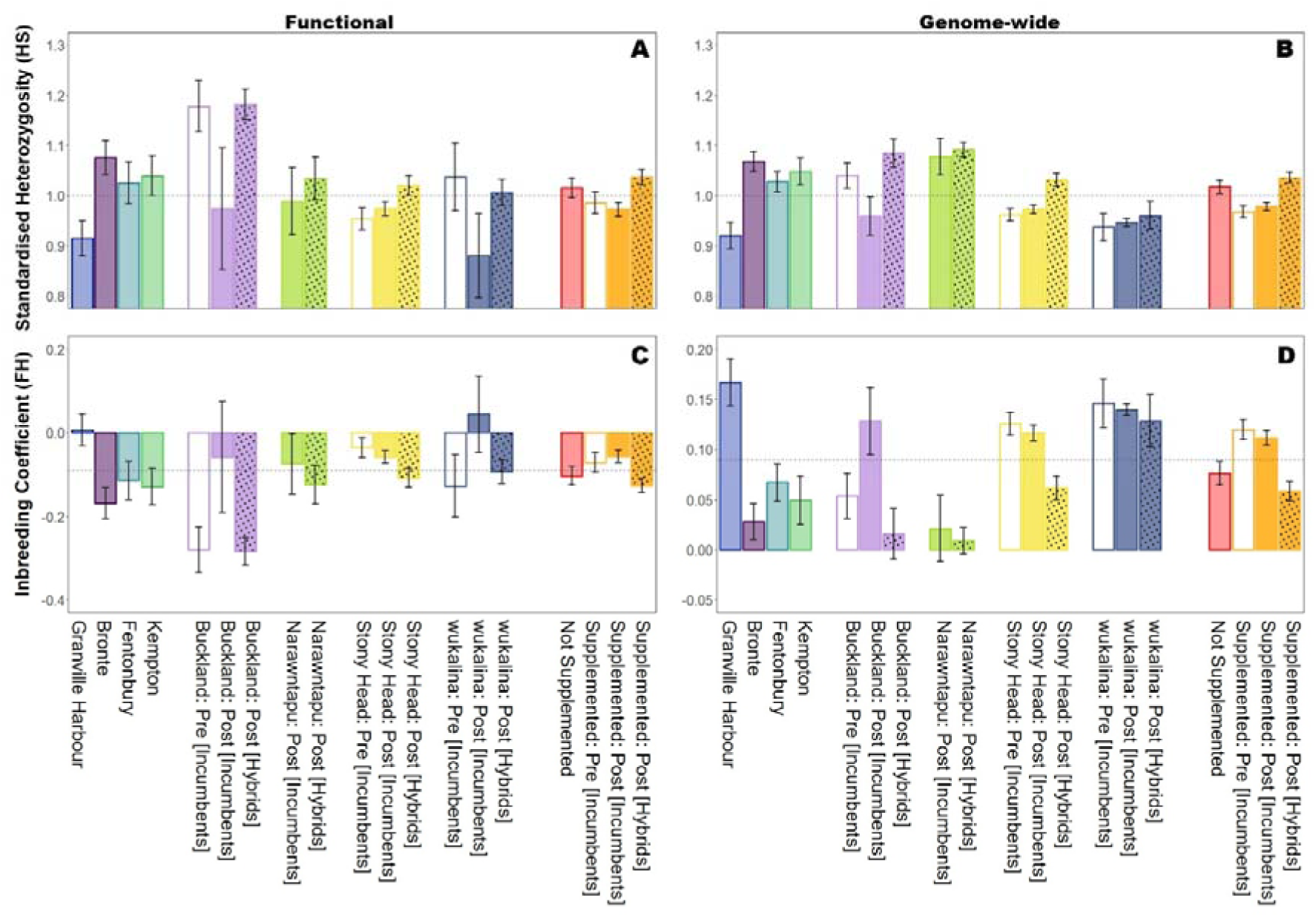
Changes in functional and genome-wide genetic diversity at not supplemented sites and at supplemented sites (pre release: incumbents only (clear bars); post release: incumbents (solid bars) and hybrids (solid bars with dots)). Standardised heterozygosity (H_S_) measured for (A) reproductive and immune-related SNPs; and (B) genome-wide SNPs. Inbreeding coefficient (FH) measured for (C) reproductive and immune-related SNPs; and (D) genome-wide SNPs. Error bars represent standard errors. Horizontal dashed lines represent the average diversity for all individuals in the dataset (reproductive and immune-related SNPs N = 325 from 379 devils; genome-wide SNPs N = 1778 from 372 devils). Sample sizes for each dataset can be found in Table S1.

Across all supplemented sites, H_S_ was either higher or the same in hybrids post supplementation compared to pre release incumbents for both functional and genome-wide diversity (Figures 2A and 2B). There appears to be no inbreeding (FH < 0) at functional genes across study sites (except Granville Harbour and post release wukalina incumbents) (Figure 2C). However, FH is accumulating at all sites when assessed with genome-wide SNPs (Figure 2D).

Across 39 reproductive and 52 immune genes (excluding MHC-I class genes), 39 haplotypes (N = 21 reproductive; N = 18 immune) were present in only hybrid individuals (Buckland, N = 7, Narawntapu, N = 16; Stony Head, N = 4; wukalina, N = 22; see supplementary Figures S1 – S8). The allelic frequencies of introduced haplotypes in supplemented sites ranged from 0.007 (CAMP_8 unique to Stony Head hybrids) to 0.350 (ADAMTS9_4 unique to Buckland hybrids). All haplotypes unique to hybrids at the supplemented sites were also detected at the not supplemented sites. There were two exceptions to this in the reproductive genes (gene name_haplotype): ADAMTS9_8 and DZIP1_6 were introduced to Stony Head (hybrid frequencies = 0.022 and 0.015, respectively). One haplotype, CD93_2, was also unique to Stony Head hybrids (frequency = 0.022), was also only seen at Granville Harbour, a not supplemented site (frequency = 0.132). The alternate haplotype, CD93_1, was fixed across all other study sites. We identified 60 haplotypes that were present in incumbents pre supplementation but absent in incumbents post supplementation. Of these, 36 haplotypes were re-introduced through the supplementation action (Buckland, N = 15, Stony Head, N = 1; wukalina, N = 20; see supplementary Figures S1 – S8), and 24 were not (Buckland, N =12, wukalina, N = 12; see supplementary Figures S1 – S8).

### MHC-I

We identified a total of 43 different MHC-I alleles (Saha-UA, N = 14; Saha-UB, N = 21; Saha-UC, N = 8) across our study sites. Overall, supplementations resulted in a slight increase in the number of MHC-I alleles per individual at supplemented sites, though we observed variability among sites (Figure 3). For not supplemented sites Granville Harbour had the highest number of MHC-I alleles, while Bronte, Fentonbury, and Kempton were all below average. Post supplementation incumbents at Buckland had a substantial decrease in the average number of MHC-I alleles. In contrast, incumbents post supplementation at both Stony Head and wukalina had more MHC-I alleles. Hybrids at Buckland and Narawntapu, two sites that had multiple supplementations events (10 – 15 individuals released every 2 years; see Schraven et al. (2025)), showed the greatest increase in MHC-I alleles compared to Stony Head and wukalina that were both “one-off” release events.

**Figure 3.**
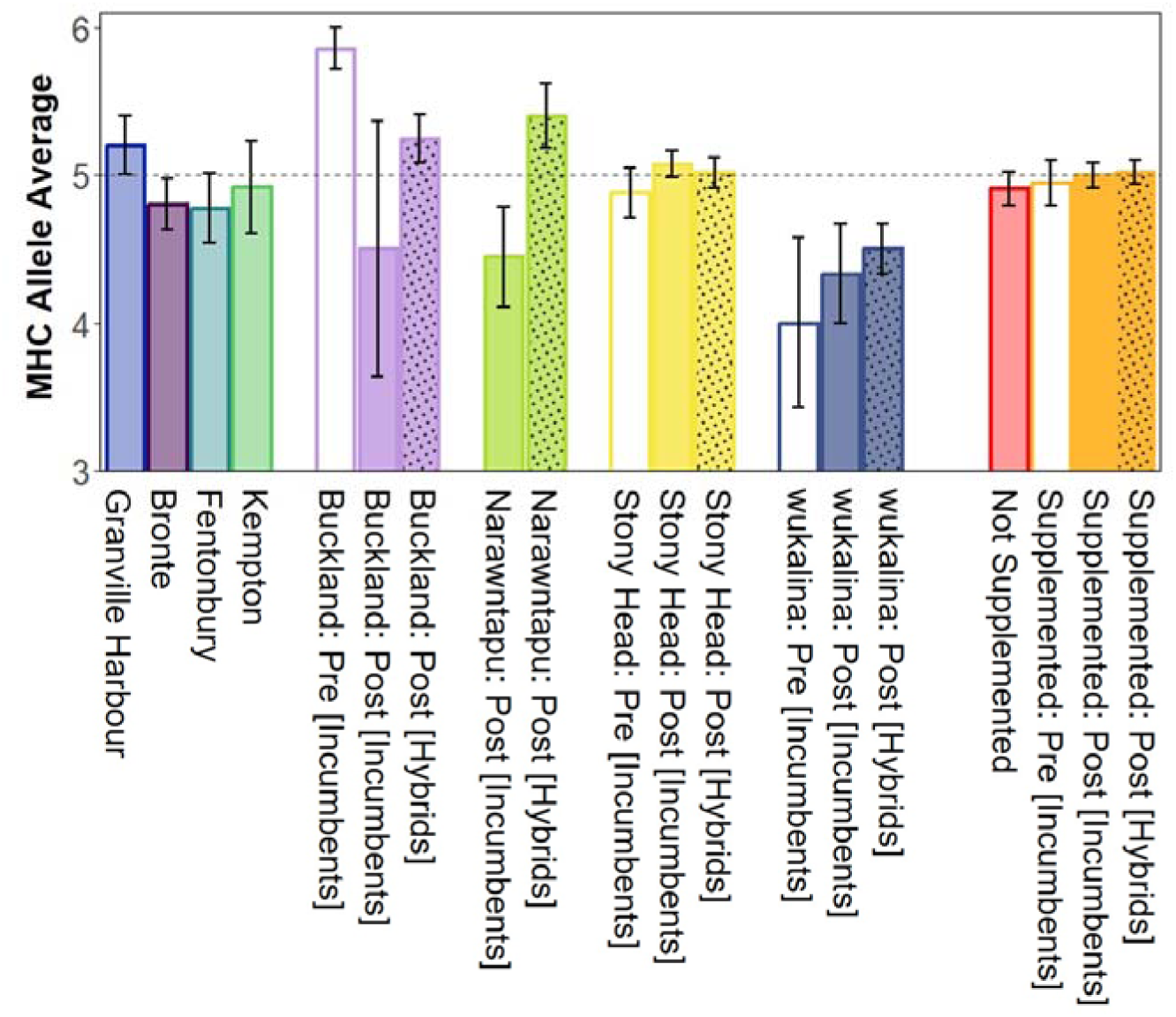
Average number of MHC alleles present in not supplemented sites and supplemented sites (pre release: incumbents only (clear bars); post release: incumbents (solid bars) and hybrids (solid bars with dots)). Error bars represent standard errors. Horizontal dashed lines represent the average number of MHC alleles present in the dataset (N = 346 devils). Sample sizes for each dataset can be found in Table S1.

We identified several MHC-I alleles unique to hybrids at Narawntapu (UA, N = 2; UB, N = 3; UC, N = 1), Stony Head (UB, N = 1), and wukalina (UA, N = 3; UB, N = 4; UC, N = 1; see Supplementary Figures S9 – S11). There were no unique MHC-I alleles in hybrids at Buckland. All these unique MHC-I alleles in hybrids were also detected in at least one of the not supplemented sites. Two of these MHC-I alleles unique to hybrids in Narawntapu (UA*01:01:01:01 frequency in hybrids = 0.05) and wukalina (UB*03:04:01:01 frequency in hybrid individuals = 0.05), were also only detected at Granville Harbour (UA and UB frequencies = 0.100 and 0.133, respectively). We observed two MHC-I UA alleles present at Buckland (UA*02:01:01:04, frequency in incumbents pre supplementation = 0.143) and wukalina (UA*01:02:02:01, frequency in incumbents pre supplementation = 0.167) pre supplementation but neither were detected post supplementation at these sites (see Supplementary Figures S9 – S11). Notably, both UA alleles were also absent from all other study sites.

Complete deletions of the UA, UB, and UC loci were variable across all study sites, with complete UA deletions represented in 16% of the dataset, complete UB at 2%, and no complete UC deletions. We also identified six individuals with both a complete UA and UB deletion (N = 5 in Stony Head and N = 1 in Buckland; all of these individuals were incumbents). Partial deletions of UA, UB, and UC were common than complete deletions, with partial UA deletion represented in 47% of the dataset, partial UB at 17%, and partial UC at 1%. At not supplemented sites, Fentonbury had the highest percentage of individuals with a complete UA deletion (39% of individuals), while Granville Harbour had the lowest percentage of individuals with a complete UA deletion (13% of individuals) (Table 1). No individuals in not supplemented sites had complete UB or UC gene deletions. No hybrid individuals in Buckland or Narawntapu (supplemented sites with multiple releases) had complete MHC I gene deletions. In contrast, hybrids in Stony Head and wukalina (both sites supplemented with a single release) had a complete UA deletion (15% and 40% of hybrids, respectively; Table 1). Only incumbents at Buckland, Narawntapu, and Stony Head had complete UB deletions (25% of incumbents in Buckland post release, 11% in Narawntapu post release, 4% in Stony Head pre release, and 2% in Stony Head post release; Table 1).

**Table 1.** Percentage of individuals within each cohort with a complete MHC I gene deletion of the UA, UB, and UC (before the “|”) d a partial deletion of the UA, UB, UC (after the “|”). Sample size for individuals genotyped for MHC I can be found in Table S1.

| Site Type | Site | MHC UA Deletions |  | MHC UB Deletions |  | MHC UC Deletions |  |
| --- | --- | --- | --- | --- | --- | --- | --- |
|  |  | Complete | Partial | Complete | Partial | Complete | Partial |
| Not Supplemented | Bronte | 20% | 60% | 0% | 20% | 0% | 0% |
|  | Fentonbury | 39% | 33% | 0% | 11% | 0% | 0% |
|  | Granville Harbour | 13% | 40% | 0% | 7% | 0% | 7% |
|  | Kempton | 33% | 17% | 0% | 17% | 0% | 8% |
| Supplemented | Buckland: Pre (Incumbents) | 0% | 14% | 0% | 0% | 0% | 0% |
|  | Buckland: Post (Incumbents) | 25% | 50% | 25% | 0% | 0% | 0% |
|  | Buckland: Post (Hybrid) | 0% | 75% | 0% | 0% | 0% | 0% |
|  | Narawntapu: Post (Incumbents) | 11% | 44% | 11% | 44% | 0% | 22% |
|  | Narawntapu: Post (Hybrids) | 0% | 40% | 0% | 10% | 0% | 10% |
|  | Stony Head: Pre (Incumbents) | 19% | 40% | 4% | 32% | 0% | 0% |
|  | Stony Head: Post (Incumbents) | 11% | 46% | 2% | 16% | 0% | 0% |
|  | Stony Head: Post (Hybrids) | 15% | 57% | 0% | 11% | 0% | 0% |
|  | wukalina: Pre (Incumbents) | 33% | 67% | 0% | 67% | 0% | 0% |
|  | wukalina: Post (Incumbents) | 33% | 67% | 0% | 33% | 0% | 0% |
|  | wukalina: Post (Hybrids) | 40% | 60% | 0% | 10% | 0% | 0% |

### DFTD

DFTD prevalence appeared to fluctuate annually among our study sites with supplementations having no impact on prevalence over time (Figure 4). DFTD was first observed at Granville Harbour in 2015 and has since shown a constant increase in prevalence over time, starting at 15**%** in 2015 to 34**%** in 2022 across individuals (15**%** to 65**%** when excluding juveniles; Figures 4A and 4C). Wild devil sites (both supplemented and not supplemented) located on the east of Tasmania show relatively similar patterns of DFTD prevalence over time (Figure 4).

**Figure 4.**
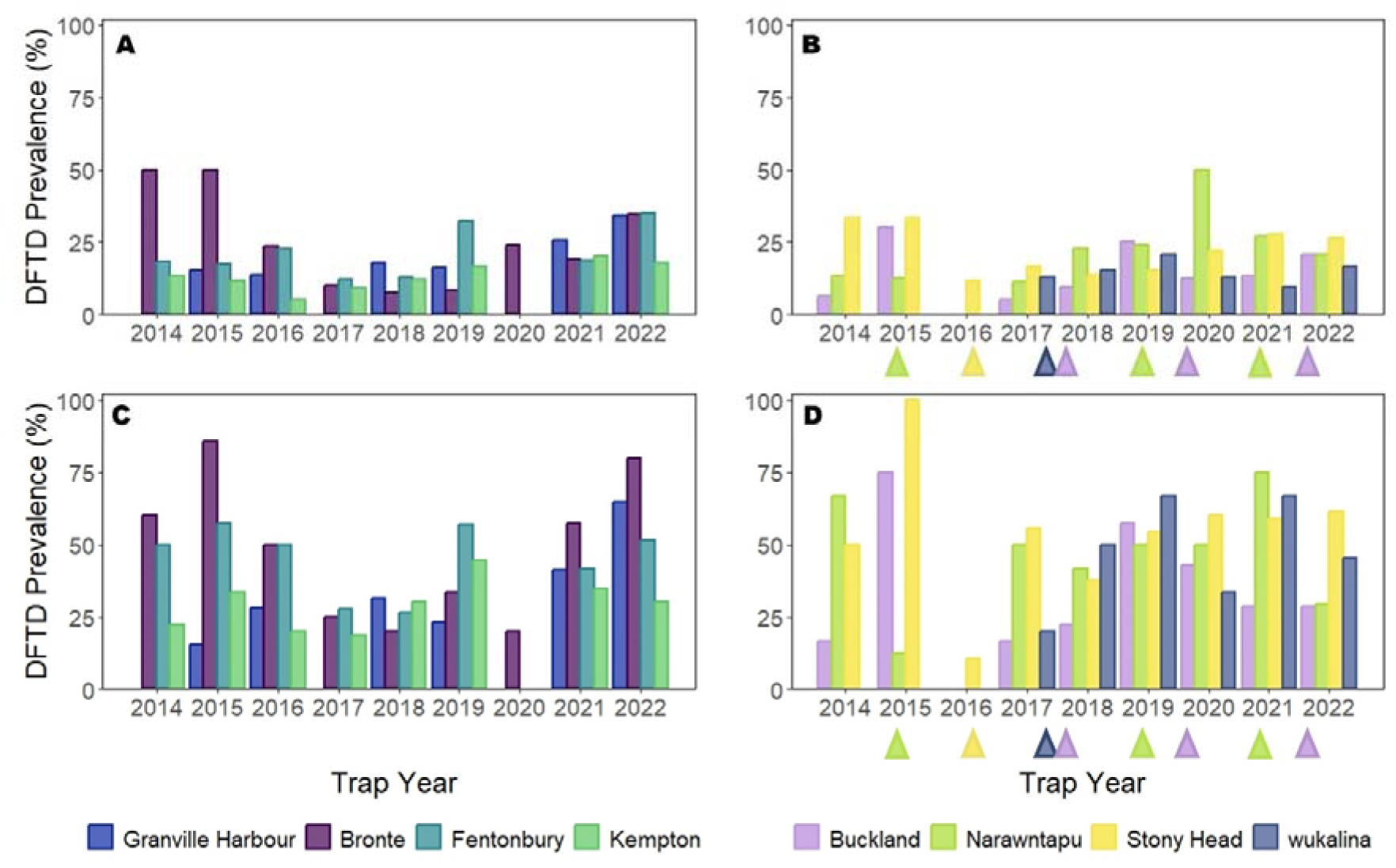
DFTD prevalence (percentage of affected devils) across trap years for (A) not supplemented and (B) supplemented sites. (C) As per (A) but excluding 1 year old devils at the time of trapping. (D) As per (B) but excluding 1 year old devils at the time of trapping. Triangles (▲) indicate which trap year devils were released into supplemented sites in (B) and (D). Note: Granville Harbour, Fentonbury, and Kempton were not trapped in 2020 due to COVID-19 related restrictions, and Granville Harbour was also not trapped in 2017.

We examined all 382 loci that have been previously reported to be under possible DFTD-driven selection (Epstein et al., 2016, Wright et al., 2017, Margres et al., 2018a). The positions of these loci on the new chromosome-level genome (Stammnitz et al., 2023) were classified each as either intergenic (N = 194), intronic (N = 166), exonic (synonymous, N = 3; nonsynonymous, N = 0), an untranslated region (UTR, N = 3), or within 500 kb upstream (N = 7) or within 500 kb downstream (N = 7) of an annotated gene. Of these 382 putative DFTD-associated loci, we identified 18 loci where variation was introduced into supplemented sites (Buckland, N = 15, Narawntapu, N = 12; see supplementary Figure S12), and three loci that were fixed post supplementation (Buckland, N = 1; Stony Head, N = 1; wukalina, N = 1; see Supplementary Figure S12). Of the three exonic loci examined, only LOC100916115 (a platelet glycoprotein VI-like gene or GP6; genomic location NC_045428.1 600198796; Supplementary Figure S12), showed a substantial allele frequency shift at supplemented sites. Variation for this locus was present at all not supplemented sites and in incumbents at Stony Head. However, the locus was fixed in incumbents for allele “C” (classified as allele_2 in our dataset) at Buckland, Narawntapu, and wukalina. Variation for this locus was introduced into Buckland and Narawntapu post supplementation, allele “T” (classified as allele_1 in our dataset) was observed in hybrids at frequencies of 0.2 and 0.05, respectively (see supplementary Figure S12). wukalina remained fixed post supplementation for this locus.

## Discussion

The wild Tasmanian devil population offers a unique study system to investigate the outcomes of management strategies that are targeted to improve adaptive potential when there is a strong selective pressure like disease. We used a targeted sequencing approach to evaluate how supplementation impacts functional diversity across multiple wild devil sites, comparing both population-level and individual-level changes pre and post supplementation. We also included genome-wide SNP data to contextualise observed patterns of functional diversity to a broader genomic scale.

Our study provides empirical evidence that multiple supplementation events have assisted gene flow of several functional alleles across eastern Tasmania, which were previously restricted to the northwestern region. Supplementations also had no impact on disease prevalence at multiple DFTD-affected sites. Our functional analysis reaffirms that devils from the insurance metapopulation are ideal candidates for supplementing wild sites (Hogg et al., 2015, 2017, 2020, Farquharson et al., 2022), and claims of their naivety toward the current selective pressure of disease are unsupported (Hohenlohe et al., 2019, Hamede et al., 2021). In the absence of wild-to-wild translocations at this time, animals for wild devil site supplementations are currently sourced from Maria Island (Hogg et al., 2020), and could also be sourced from Forestier Peninsula (Huxtable et al., 2019). From previous landscape-level analyses we know that free-living disease free sites established from the insurance population genetically represent wild genome-wide and functional diversity (Farquharson et al., 2022). In addition, Maria Island is actively managed to ensure genetic diversity is maximised long-term (Hogg et al., 2020).

Supplementations appeared to contribute to the reduction in genetic differentiation between sites, while also increasing both population- and individual-level functional diversity and reducing inbreeding. Our previous investigation into genome-wide diversity showed that supplementations had a site-specific impact on diversity over time with three supplemented sites increasing (Narawntapu, Stony Head, and wukalina) and one supplemented site (Buckland) decreasing over time (Schraven et al., 2025). By comparing incumbents pre and post supplementation to hybrids (post supplementation), we observed similar changes in both functional and genome-wide diversity pre and post supplementation. Interestingly, hybrids at Buckland had similar levels of functional and genome-wide diversity and inbreeding to incumbents pre supplementations. However, our reconstructed pedigrees of supplemented sites showed that the number of hybrids present at Buckland is small compared to other supplemented sites. This suggests that individuals released at Buckland may disperse from the local area prior to mating with incumbents, resulting in no measurable impact on the population-level variation. This also aligns with our current knowledge of gene flow in this area, whereby Buckland has a higher level of gene flow than many other sites (Schraven et al., 2024).

We identified 52 functional haplotypes that were introduced across the four supplemented wild devil sites. Of these, all but two (ADAMTS9_8 and DZIP1_6) were already present within the broader landscape, indicating that supplementation efforts have primarily acted to redistribute existing genetic variation rather than introducing entirely novel alleles. This supports the current management practices of prioritising release sites to coastal and relatively isolated devil sites (Grueber et al., 2019, Schraven et al., 2024). Assisted gene flow of functional genes has been shown to increase the adaptive potential of threatened populations in rapidly changing environments (Kelly and Phillips, 2016). Where natural gene flow is limited, between the east and western located devil sites for example (Schraven et al., 2024), assisted gene flow may facilitate the increase of evolutionary resilience (Kelly and Phillips, 2016). For example, prior to supplementation, there was no variation of the immune CD93 gene across eastern Tasmania, as any variation in this locus was restricted to the northwest region. This northwest region variation was subsequently introduced to Stony Head through supplementation efforts (McLennan et al., 2024). The two newly introduced reproductive haplotypes, ADAMTS9_8 and DZIP1_6, have been previously characterized in the devil (Brandies et al., 2021). ADAMTS9 is a member of the ADAMTS protease family, involved in multiple female reproductive processes, such as ovulation, implantation, and placentation (Brandies et al., 2021). ADAMTS is also known to have tumour suppressor functionality for multiple cancers (Latifi et al., 2014). DZIP1 is a regulator of hedgehog signalling (Schwend et al., 2013), though recent bioinformatic analysis has indicated potential associations between expression levels of DZIP1 and survival in human cancers (Liu et al., 2021). It is also noted that all 52 haplotypes were introduced at relatively low frequencies. This reduces the likelihood of overwhelming locally adapted gene pools while still enhancing overall functional genetic diversity.

In addition to the influx of immune-related and reproductive diversity in supplemented sites, we found that supplementations also increased the number of MHC-I alleles present. Increased MHC-I diversity in a species that has known low immune gene diversity provides a pathway to respond to disease (Morris et al., 2013, Cheng et al., 2017). MHC-I genes have been implicated in the spread of DFT1, whereby the tumour evades the host’s immune system by downregulating MHC-I on their cell surface (Siddle et al., 2013). Extremely low levels of MHC-I diversity, much of which is shared with DFT1 cells, means that DFT1 can hide from the devil’s immune system (Morris et al., 2013). Recently we have observed that individuals with either a complete gene deletion of the MHC-I UA gene are more likely to produce antibodies against DFT1 cells (Batley et al., 2025). The complete UA deletion is possibly being selected for at long term disease sites, however, it does not necessarily result in individual survivability following DFT1 infection (Batley et al., 2025). If an increase in the frequency of deletion at UA reflects an evolutionary response to DFT1, our results suggest that our supplementation activities have not acted counter to natural processes as our supplementations increased the frequency of the UA deletion. This is because many of the founders from the insurance population were predominantly from the west coast where there is a known higher frequency of the UA deletion (Cheng et al., 2012, Lane et al., 2012, Hogg et al., 2015).

Supplementing threatened populations in the presence of infectious diseases can be problematic. However, as DFT1 is a frequency-dependent disease (McCallum et al., 2009), the influx of new individuals does not impact disease prevalence. Rather DFT1 prevalence naturally fluctuates on a temporal scale across all our study sites, regardless of if they were supplemented or not supplemented. While considerable effort has been invested into a vaccine for DFTD (Pye et al., 2018, Pye et al., 2021), an efficient and deployable solution is still in development. Therefore, the current management strategy of the STDP continues to focus on monitoring wild devil populations in the presence of DFTD. Here we also explored specific regions of the devil genome that had previously been implicated in potential tolerance and/or resistance to DFT1 (Epstein et al., 2016, Wright et al., 2017, Margres et al., 2018a). Using the new scaffolded-assembly genome, we show that most of these regions do not reside within exonic regions of a gene. Only three of these putative DFTD-associated loci are located within exons, and none result in an amino acid change. Of these three loci, we only introduced variation of one of these loci (LOC100916115 or GP6) at two supplemented sites (Buckland and Narawntapu). The variation we introduced at these sites are already present at all not supplemented sites. This suggests that supplementations have assisted gene flow of current standing genetic variation rather than introducing novel adaptive alleles. However, further functional validation is needed to assess whether these loci play a causal role in DFT1 tolerance and/or resistance.

These findings need to be considered within the broader context of uncertainty around the genetic basis of the devil’s response to DFTD and future potential threats. In small and isolated populations, such as those found across the devil’s natural range, genetic drift can outweigh any potential for natural selection of advantageous genotypes (Bouzat, 2010). For instance, in a reintroduced population of Stewart Island robins (*Petroica australis rakiura*), individuals with the TLR4_BE_ genotype showed improved survival, yet the allele frequency of TLR4_BE_ remained too low for natural selection to act effectively (Grueber et al., 2013). Two independent epidemiological models have suggested that devils may persist and even coexist with DFT1 (Siska et al., 2018, Wells et al., 2019), a finding that has contributed to the ongoing debate regarding long-term management strategies for devils. However, both models predicted population sizes would remain significantly low, which not only compromises the species’ demographic and genetic viability but may also increase ecological disruption. These insights, combined with the current uncertainty surrounding the natural selection for resistance towards DFTD, underscores the importance of ongoing management support for the devil at this time.

## Conclusions

To safeguard the future of species facing diminished adaptive potential under increasing selective pressures, like disease, it is necessary to understand functional genetic diversity and how it may contribute to disease resilience. We have demonstrated that supplementations successfully increased overall functional genetic diversity in Tasmanian devils, strengthening their long-term persistence in the wild. Disease as a significant threatening process is not unique to devils, nor will it be the only emerging challenge on the horizon. It is therefore vital to equip devils, and similarly vulnerable species, with species-level genetic variability needed for long-term population viability. We have shown that current supplementation strategies do not swamp existing genetic variants but instead enhance their variability across the fragmented landscape. Additionally, while our focus was the impact of supplementation on functional diversity, we also compared these changes to genome-wide diversity. Observed changes in genome-wide diversity closely mirrored those in functional diversity, supporting the use of genome-wide diversity as a proxy for adaptive potential in conservation management where sequencing functional genes may not be feasible (Kardos et al., 2021). For devils, future monitoring will be needed to determine whether the increased functional variation as a result of supplementations is maintained long-term, or if genetic drift and/or selective pressures lead to a recurrence of population differentiation and loss of genetic diversity over time.

## Acknowledgements

We acknowledge the traditional custodians of the land on which these devil (purinina) populations live and pay respects to their elders past and present. We thank the various current and past staff of the Save the Tasmanian Devil Program who have undertaken extensive monitoring of devil sites and participated in the sourcing and release of devils, in particular Jodie Elmer, David Schaap, Bill Brown, Stewart Huxtable, Clare Lawrence, and Phil Wise. This work was supported by the Australian Research Council [LP180100244], the University of Sydney, the Save the Tasmanian Devil Program and San Diego Zoo Wildlife Alliance. KF and LS are supported by funding from the ARC Centre of Excellence for Innovations in Peptide and Protein Science (CE200100012).

## Data availability

All raw sequence data will be deposited onto Dryad following acceptance of this manuscript.

## Benefit-Sharing Statement

A research collaboration exists between academics and the Department of Natural Resources and Environment Tasmania (NRE Tas), who provided operational and logistical support, including access to information and genetic samples to undertake this study. The results of the research have been shared with the NRE Tas in real-time to inform their management actions for the conservation of the Tasmanian devil. NRE Tas engage with local Indigenous communities and other local communities, particularly during the establishment of the Wild Devil Recovery Project and the first supplementations to sites.

## Author contributions

Conceptualisation, C.J.H, K.B.; Fieldwork, C.J.H., S.F., A.V.L; Data analysis, A.L.S., L.S., K.A.F., K.C.B.; Writing – Original Draft, A.L.S., L.S., K.A.F.; Writing – Review and Editing, all authors; Visualisation, A.L.S.; Supervision, C.J.H.; Resources, C.J.H., K.B., S.F.; Funding acquisition, C.J.H., S.F., K.

