## Supplementary material for "Boost or Bust? The impact of supplementation on functional genetic diversity and selective processes in Tasmanian devils"

Carolyn J. Hogg: 0000-0002-6328-398X

**This file includes:**

Table S1

Table S2

Figure S1

Figure S3

Figure S4

Figure S5

Figure S6

Figure S7

Figure S8

Figure S9

Figure S10

Figure S11

Figure S12

Table S1. Sample sizes of devils included in this study.

| Site Type | Site | Functional | MHC | Genome-wide |
| --- | --- | --- | --- | --- |
| Not Supplemented | Bronte | 21 | 20 | 21 |
|  | Fentonbury | 20 | 18 | 19 |
|  | Granville Harbour | 19 | 15 | 19 |
|  | Kempton | 20 | 12 | 20 |
| Supplemented | Buckland: Pre (Incumbents) | 7 | 7 | 7 |
|  | Buckland: Post (Incumbents) | 4 | 4 | 3 |
|  | Buckland: Post (Hybrid) | 10 | 8 | 10 |
|  | Narawntapu: Post (Incumbents) | 9 | 9 | 8 |
|  | Narawntapu: Post (Hybrids) | 10 | 10 | 10 |
|  | Stony Head: Pre (Incumbents) | 52^a^ | 47^a^ | 51^a^ |
|  | Stony Head: Post (Incumbents) | 125^a^ | 122^a^ | 124^a^ |
|  | Stony Head: Post (Hybrids) | 67 | 61 | 65 |
|  | wukalina: Pre (Incumbents) | 6 | 3 | 6 |
|  | wukalina: Post (Incumbents) | 3 | 3 | 3 |
|  | wukalina: Post (Hybrids) | 10 | 10 | 10 |

^a^four incumbent individuals were trapped at Stony Head both pre and post supplementation, these are included twice in the sample sizes.

Table S2. Pairwise F_ST_ values between wild devil sites. Below the diagonal denotes the F_ST_ values for functional diversity (excluding MHC I and DFTD) and above the diagonal denotes the F_ST_ values for genome-wide diversity. All values were significantly different from 0 at p < 0.05.

|  | Granville Harbour | Bronte | Fentonbury | Kempton | Buckland | Narawntapu | Stony Head | wukalina |
| --- | --- | --- | --- | --- | --- | --- | --- | --- |
| Granville Harbour | - | 0.095 | 0.108 | 0.091 | 0.094 | 0.123 | 0.125 | 0.128 |
| Bronte | 0.072 | - | 0.026 | 0.017 | 0.037 | 0.053 | 0.069 | 0.077 |
| Fentonbury | 0.074 | 0.012 | - | 0.017 | 0.040 | 0.062 | 0.072 | 0.079 |
| Kempton | 0.094 | 0.020 | 0.016 | - | 0.028 | 0.047 | 0.055 | 0.062 |
| Buckland | 0.094 | 0.040 | 0.045 | 0.033 | - | 0.048 | 0.068 | 0.044 |
| Narawntapu | 0.124 | 0.069 | 0.068 | 0.064 | 0.055 | - | 0.068 | 0.069 |
| Stony Head | 0.116 | 0.057 | 0.067 | 0.062 | 0.043 | 0.057 | - | 0.045 |
| wukalina | 0.111 | 0.048 | 0.052 | 0.049 | 0.042 | 0.058 | 0.082 | - |

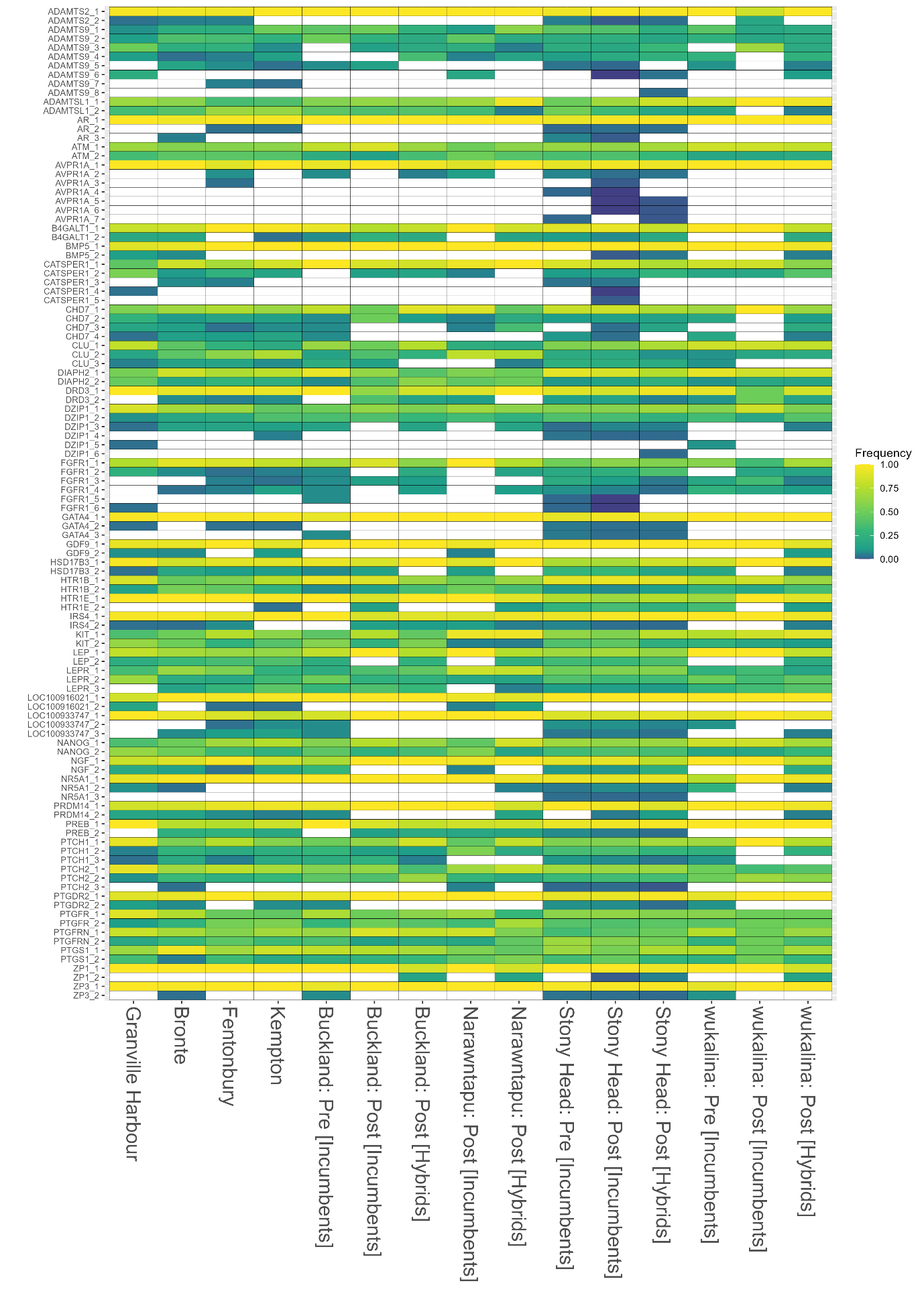

**Figure S1.** Heatmap of allele frequencies of haplotypes at coding regions of reproductive genes. Each column represents wild devil site, including not supplemented sites, supplemented sites pre supplementation (incumbents only), and supplemented sites post supplemented (incumbents and hybrids). Each row represents a haplotype where they are ordered alphabetically and shaded by their frequency corresponding to the scale bar. Rows are labelled by haplotype names as designated during phasing.

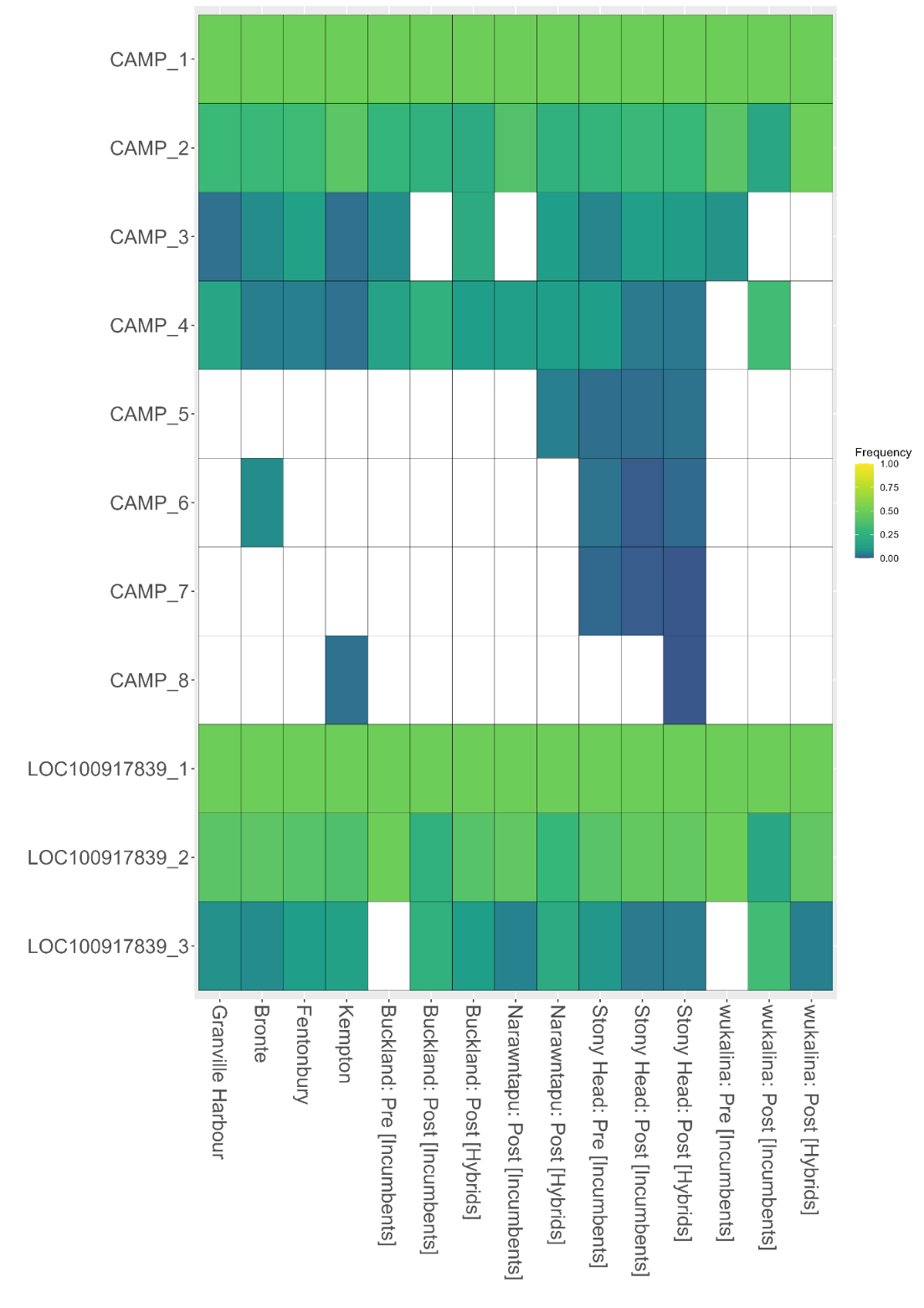

**Figure S2.** Heatmap of allele frequencies of haplotypes at coding regions of antimicrobial peptide (AMP) genes. Each column represents wild devil site, including not supplemented sites, supplemented sites pre supplementation (incumbents only), and supplemented sites post supplemented (incumbents and hybrids). Each row represents a haplotype where they are ordered alphabetically and shaded by their frequency corresponding to the scale bar. Rows are labelled by haplotype names as designated during phasing.

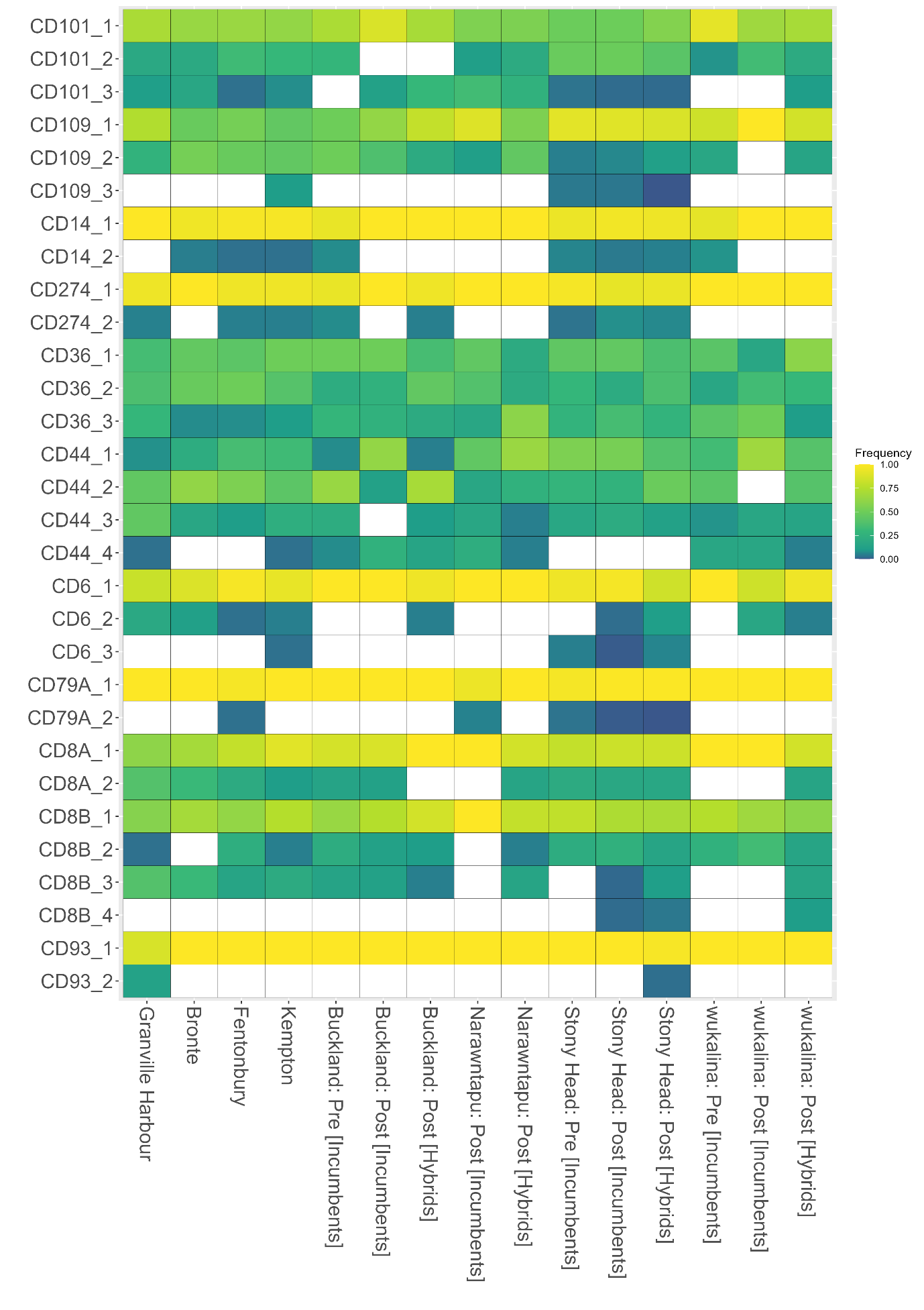

**Figure S3.** Heatmap of allele frequencies of haplotypes at coding regions of cluster of differentiation (CD) genes. Each column represents wild devil site, including not supplemented sites, supplemented sites pre supplementation (incumbents only), and supplemented sites post supplemented (incumbents and hybrids). Each row represents a haplotype where they are ordered alphabetically and shaded by their frequency corresponding to the scale bar. Rows are labelled by haplotype names as designated during phasing.

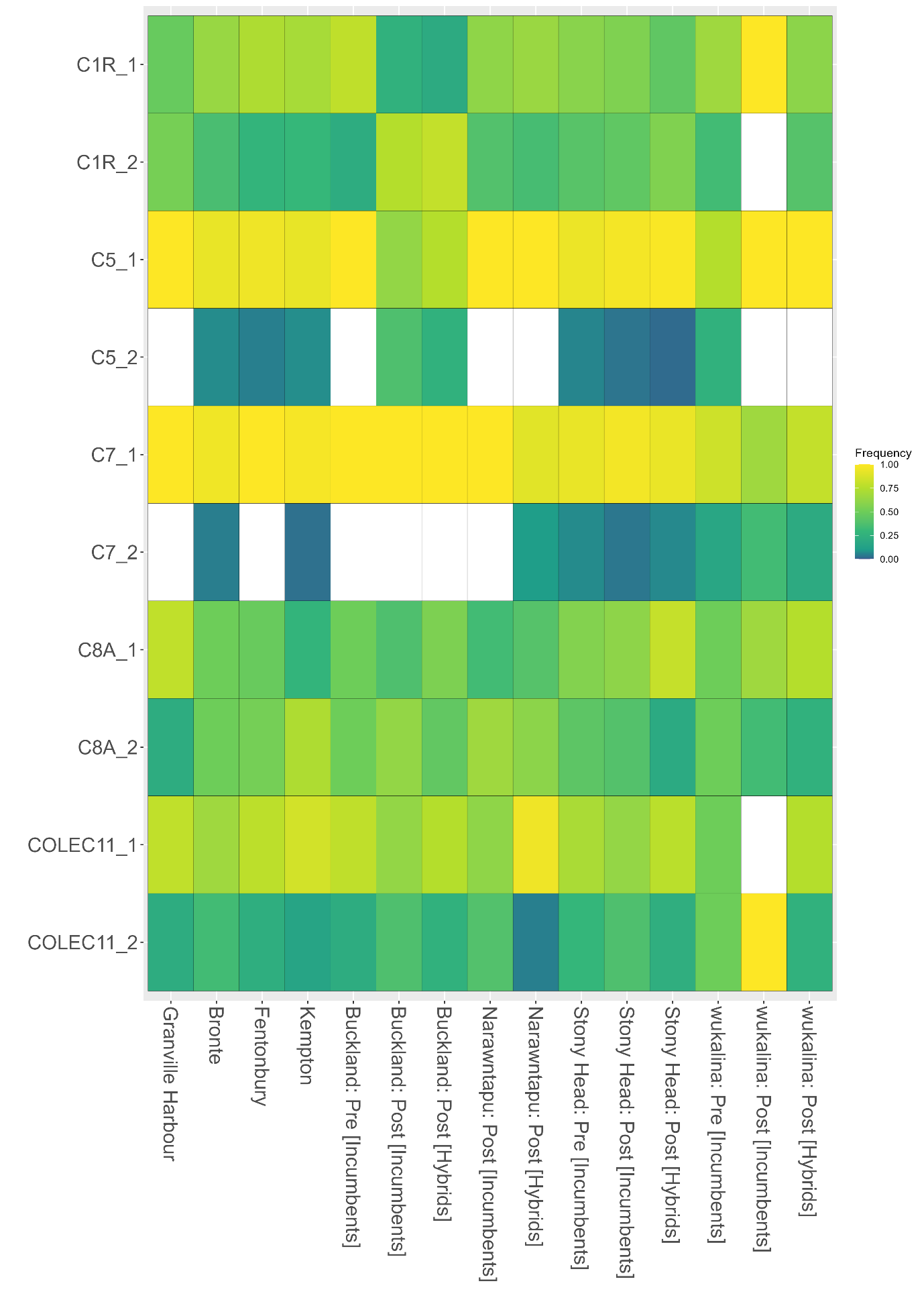

**Figure S4.** Heatmap of allele frequencies of haplotypes at coding regions of complement genes. Each column represents wild devil site, including not supplemented sites, supplemented sites pre supplementation (incumbents only), and supplemented sites post supplemented (incumbents and hybrids). Each row represents a haplotype where they are ordered alphabetically and shaded by their frequency corresponding to the scale bar. Rows are labelled by haplotype names as designated during phasing.

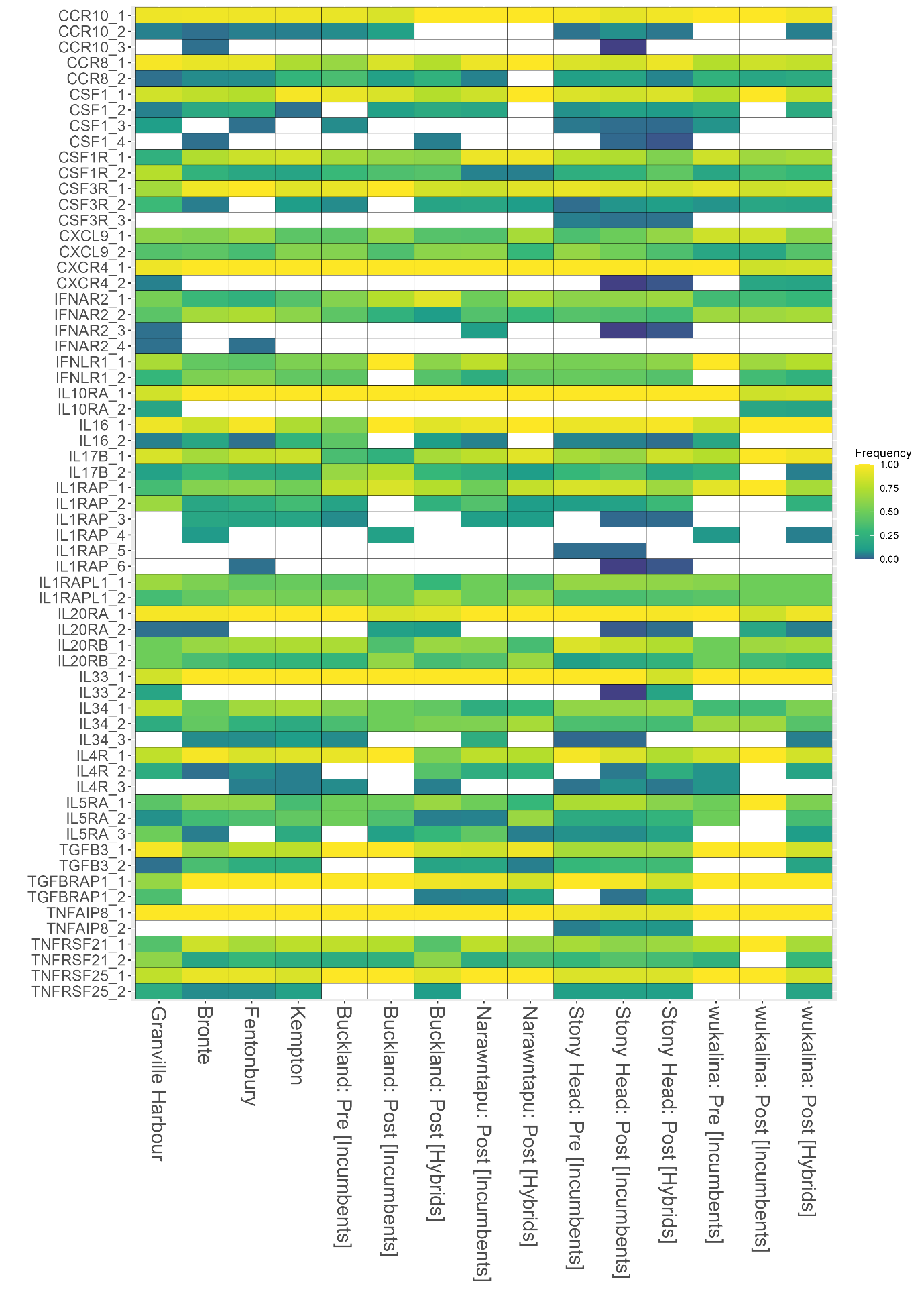

**Figure S5.** Heatmap of allele frequencies of haplotypes at coding regions of cytokine genes. Each column represents wild devil site, including not supplemented sites, supplemented sites pre supplementation (incumbents only), and supplemented sites post supplemented (incumbents and hybrids). Each row represents a haplotype where they are ordered alphabetically and shaded by their frequency corresponding to the scale bar. Rows are labelled by haplotype names as designated during phasing.

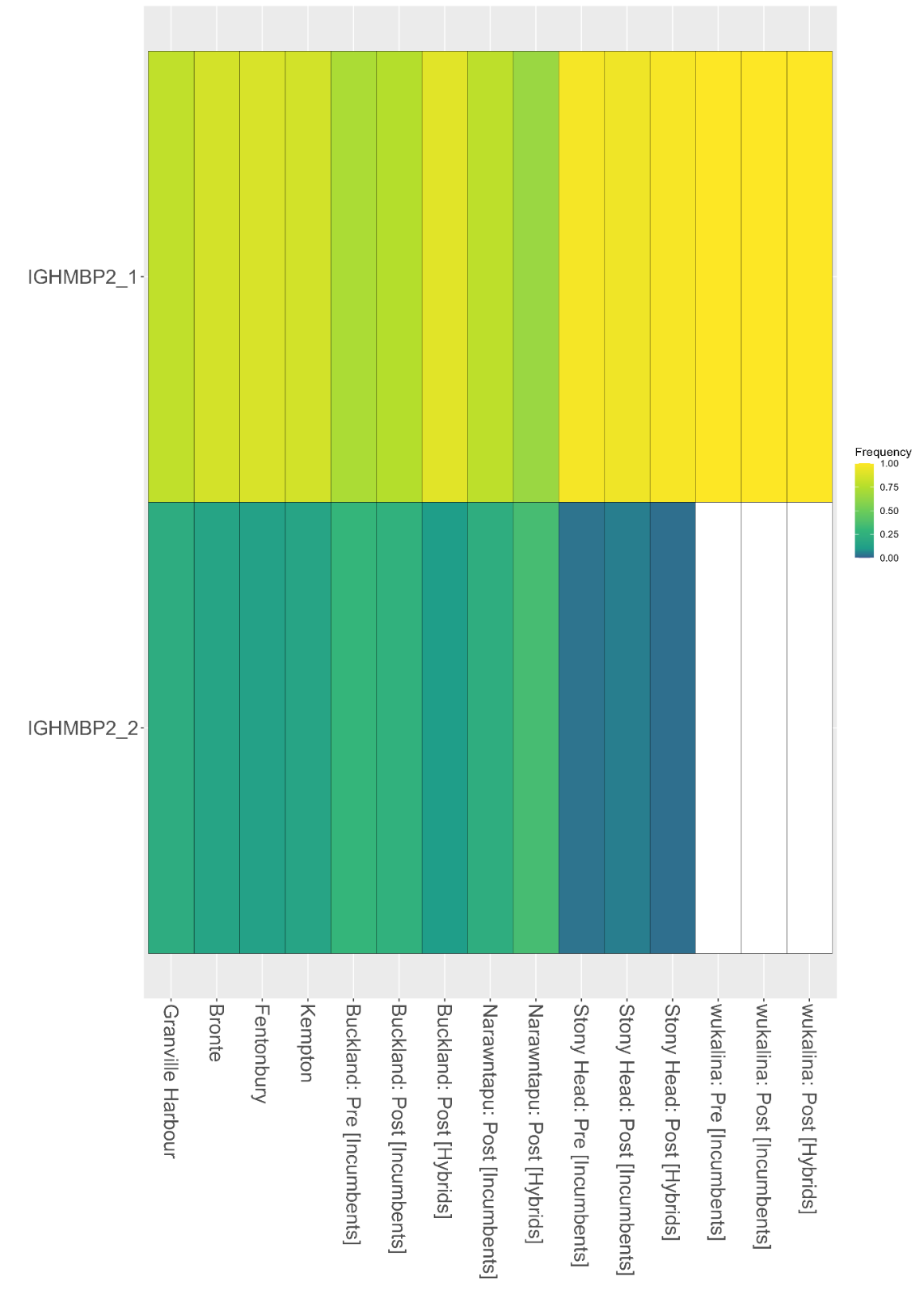

**Figure S6.** Heatmap of allele frequencies of haplotypes at coding regions of immunoglobulin genes. Each column represents wild devil site, including not supplemented sites, supplemented sites pre supplementation (incumbents only), and supplemented sites post supplemented (incumbents and hybrids). Each row represents a haplotype where they are ordered alphabetically and shaded by their frequency corresponding to the scale bar. Rows are labelled by haplotype names as designated during phasing.

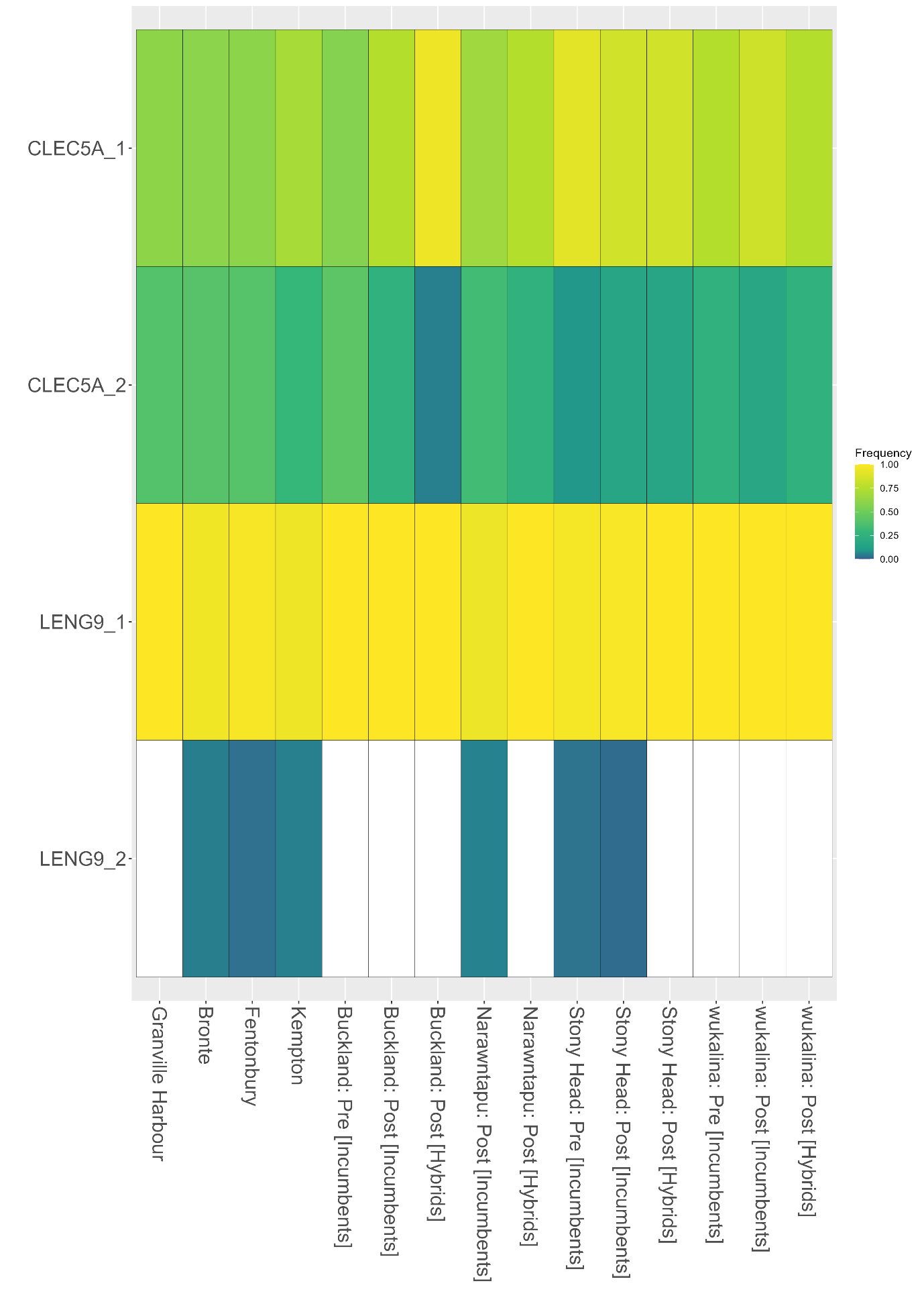

**Figure S7.** Heatmap of allele frequencies of haplotypes at coding regions of natural killer cell receptor (NK) genes. Each column represents wild devil site, including not supplemented sites, supplemented sites pre supplementation (incumbents only), and supplemented sites post supplemented (incumbents and hybrids). Each row represents a haplotype where they are ordered alphabetically and shaded by their frequency corresponding to the scale bar. Rows are labelled by haplotype names as designated during phasing.

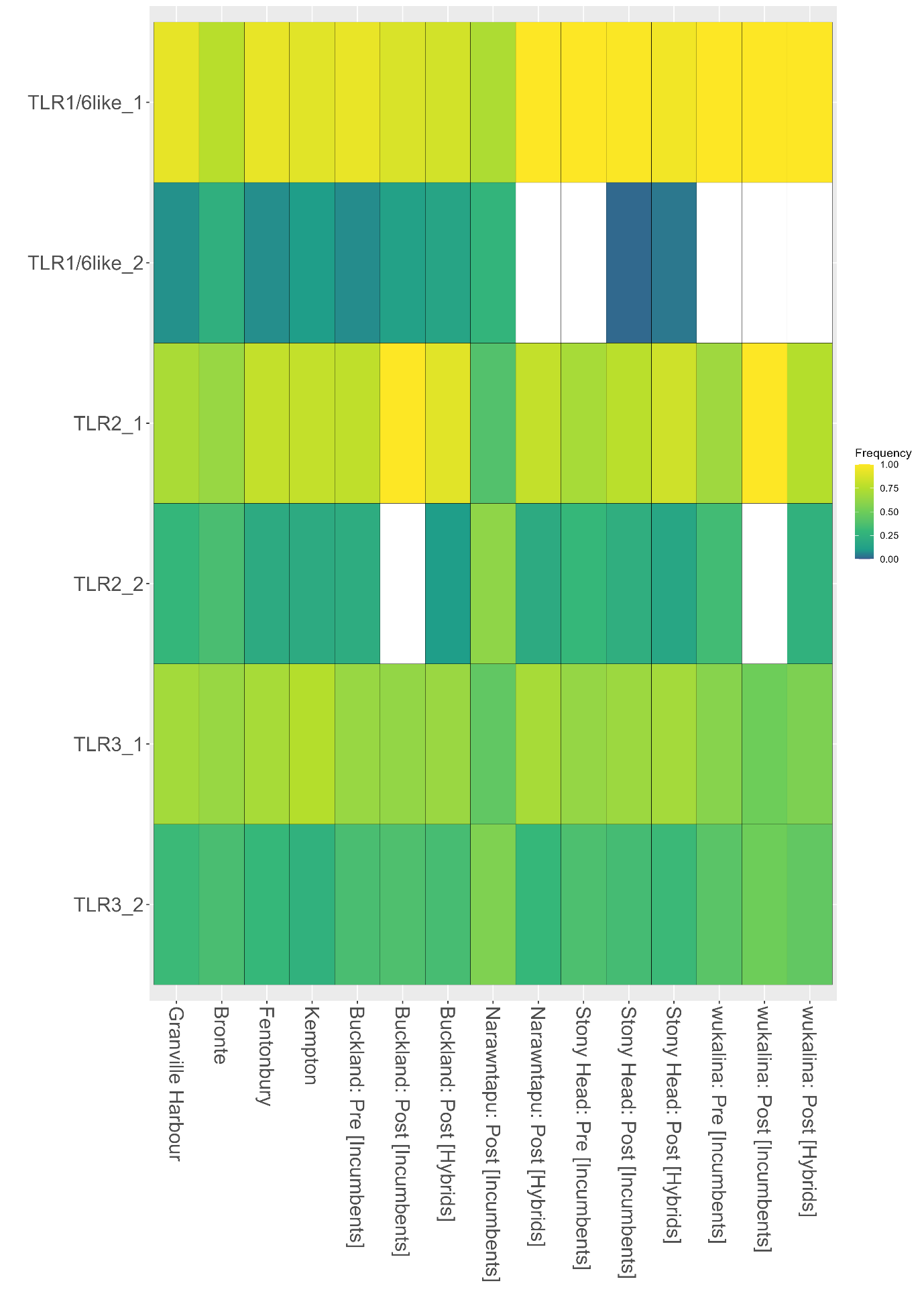

**Figure S8.** Heatmap of allele frequencies of haplotypes at coding regions of toll-like receptor (TLR) genes. Each column represents wild devil site, including not supplemented sites, supplemented sites pre supplementation (incumbents only), and supplemented sites post supplemented (incumbents and hybrids). Each row represents a haplotype where they are ordered alphabetically and shaded by their frequency corresponding to the scale bar. Rows are labelled by haplotype names as designated during phasing.

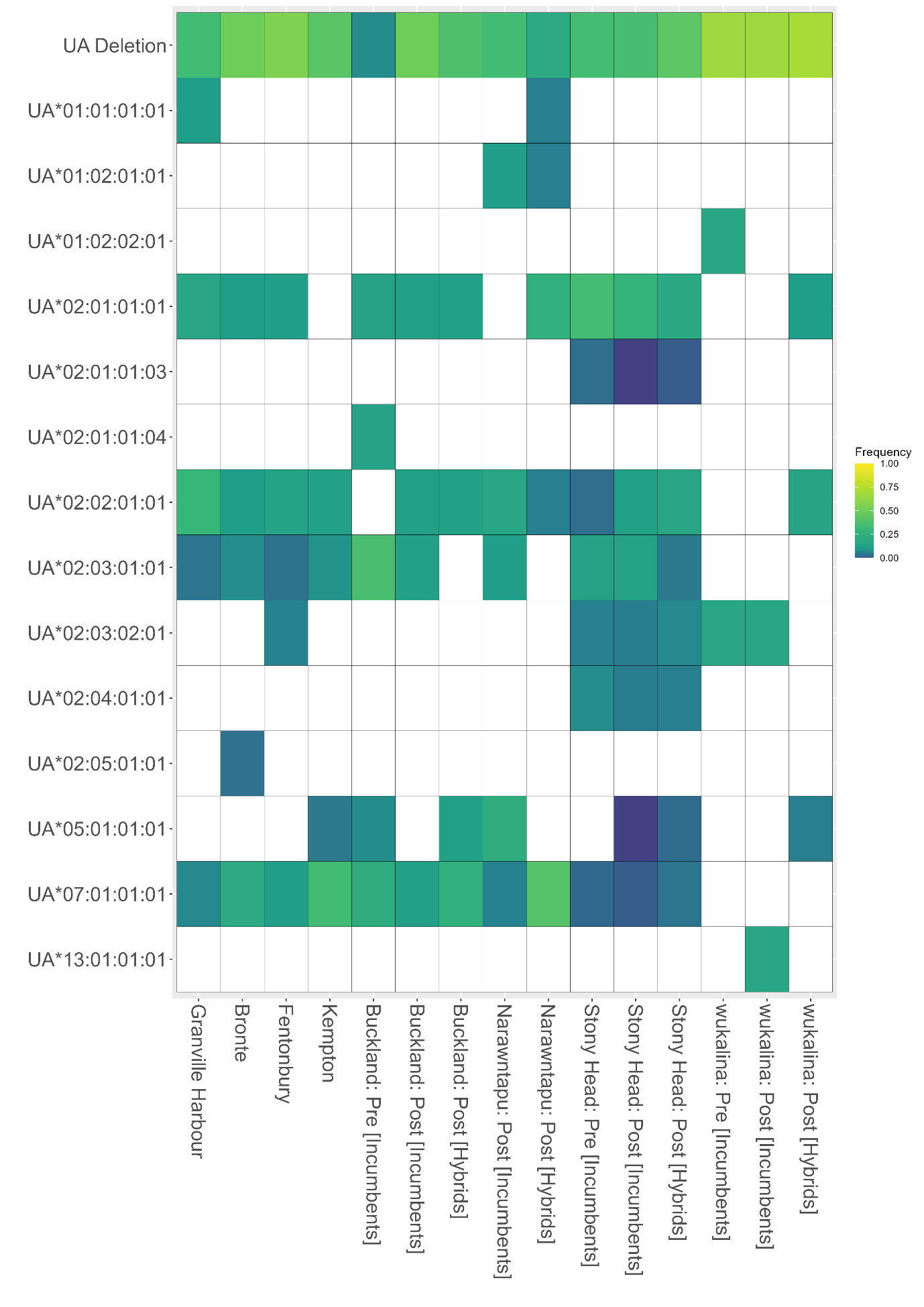

**Figure S9.** Heatmap of allele frequencies of MHC class I UA genes. Each column represents wild devil site, including not supplemented sites, supplemented sites pre supplementation (incumbents only), and supplemented sites post supplemented (incumbents and hybrids). Each row represents a UA allele, with the top row indicating a deletion of a UA allele. The frequency of each allele is shown according to the scale bar.

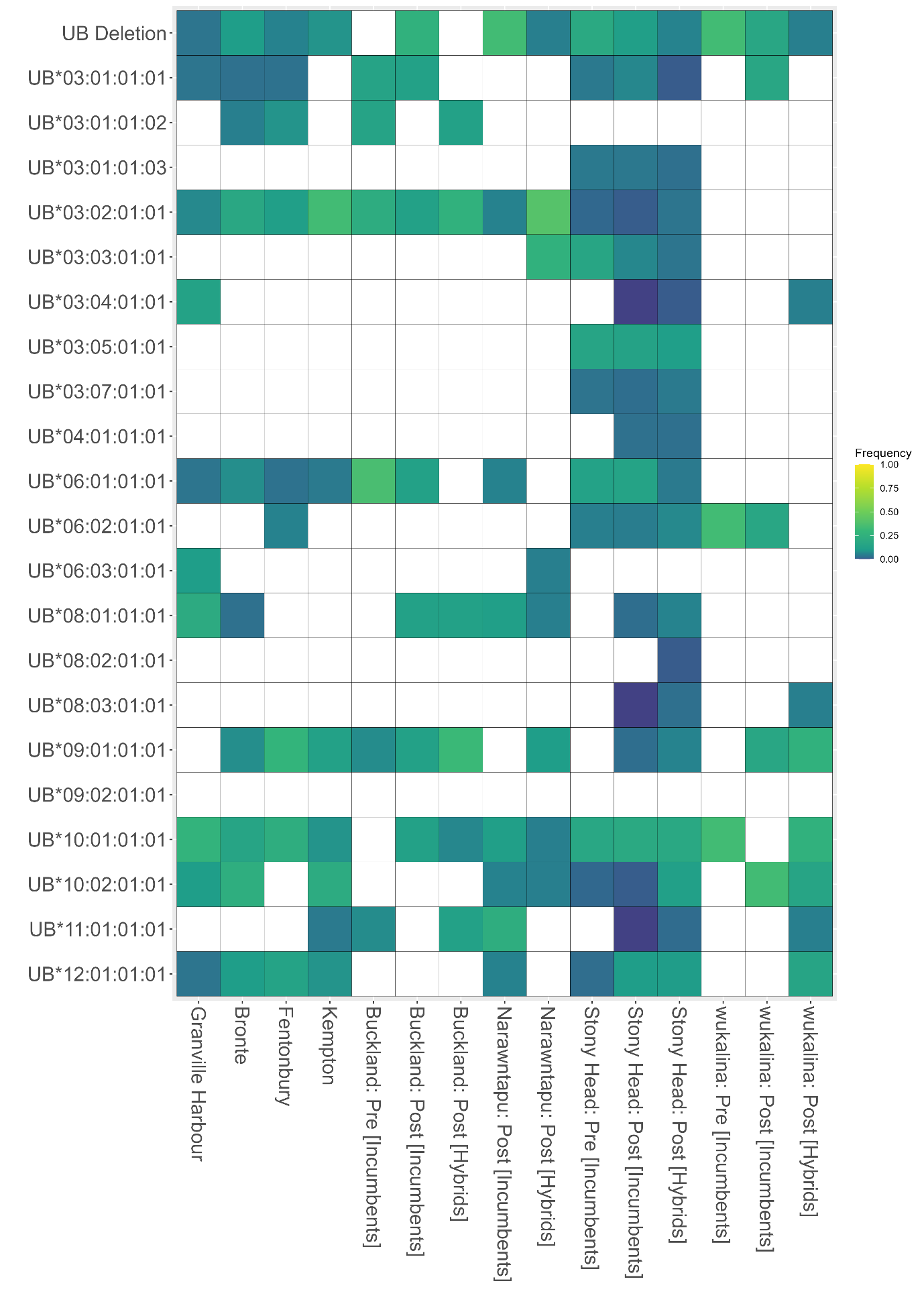

**Figure S10.** Heatmap of allele frequencies of MHC class I UB genes. Each column represents wild devil site, including not supplemented sites, supplemented sites pre supplementation (incumbents only), and supplemented sites post supplemented (incumbents and hybrids). Each row represents a UB allele, with the top row indicating a deletion of a UB allele. The frequency of each allele is shown according to the scale bar.

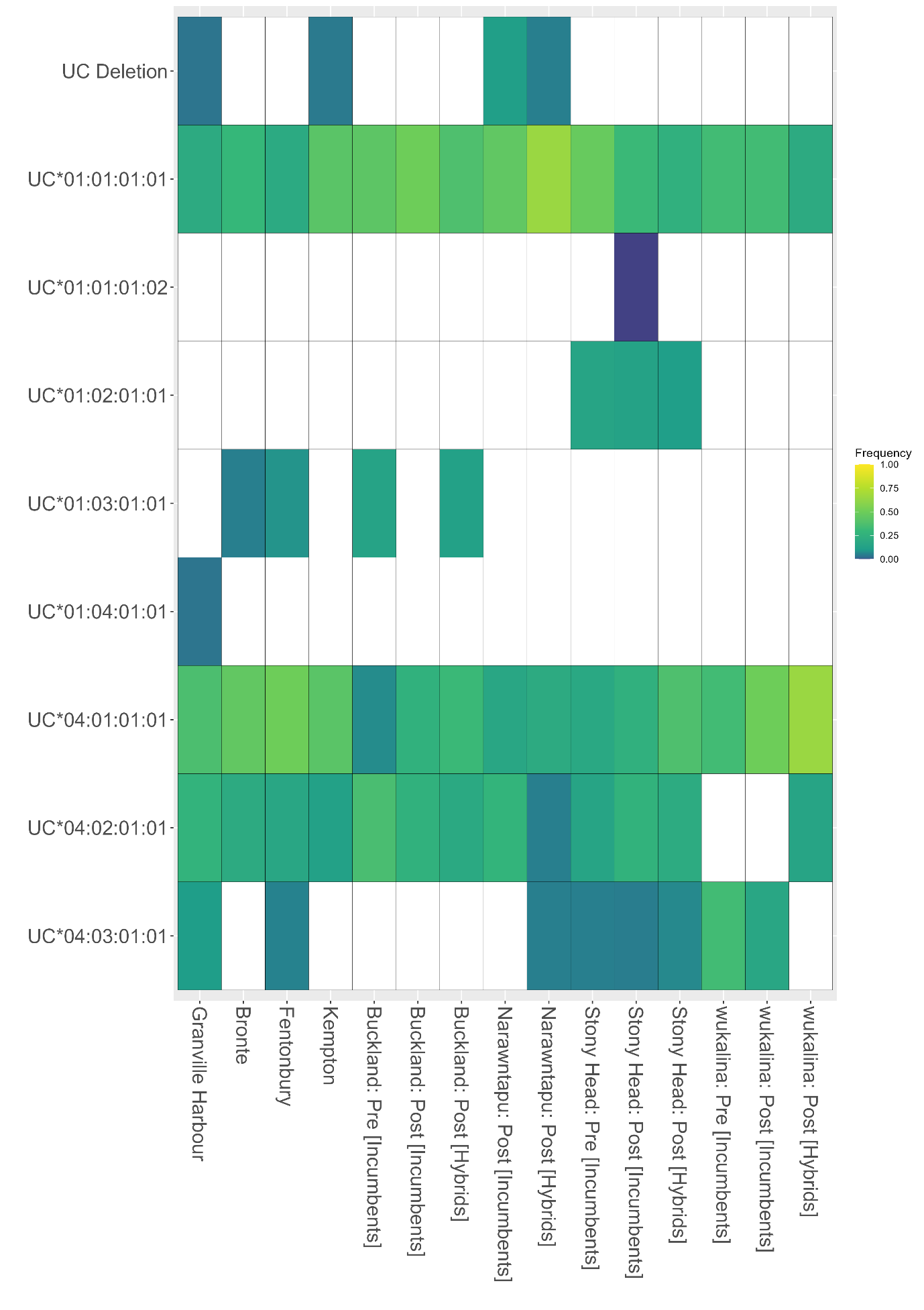

**Figure S11**. Heatmap of allele frequencies of MHC class I UC genes. Each column represents wild devil site, including not supplemented sites, supplemented sites pre supplementation (incumbents only), and supplemented sites post supplemented (incumbents and hybrids). Each row represents a UC allele, with the top row indicating a deletion of a UC allele. The frequency of each allele is shown according to the scale bar.

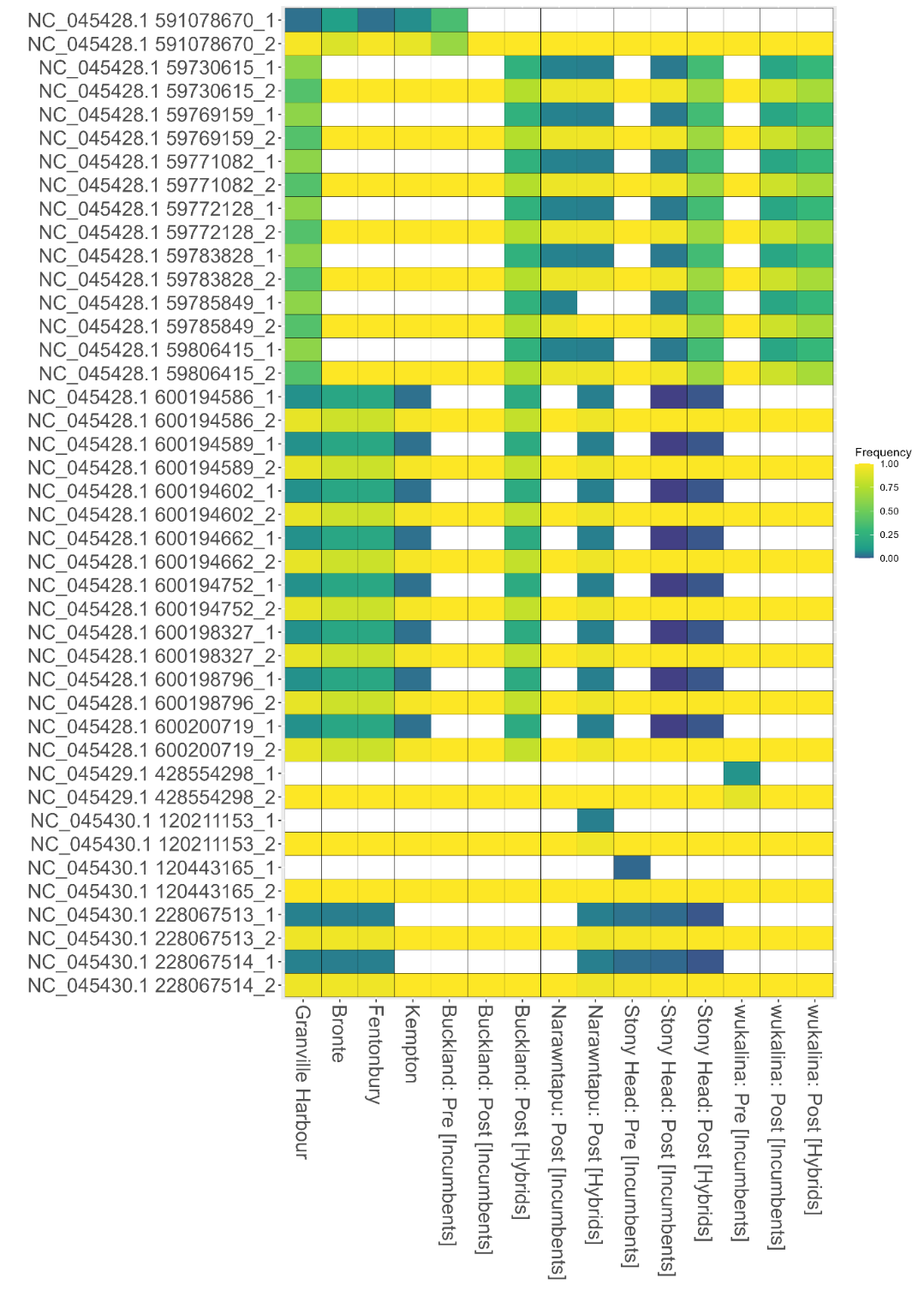

**Figure S12.** Heatmap of allele frequencies of the 21 putative DFTD-loci which were either introduced to supplemented sites (i.e. unique to hybrids post supplementation, N = 18) or were fixed at supplemented sites (i.e. unique to incumbents pre supplementation, N = 3). Each column represents wild devil site, including not supplemented sites, supplemented sites pre supplementation (incumbents only), and supplemented sites post supplemented (incumbents and hybrids). Each row represents a putative DFTD allele (labelled by their genomic position of which the REF allele is denoted by the “_1” and the ALT allele is denoted by the "_2” at the end of the label) and are shaded by their frequency corresponding to the scale bar.
